# Computational and *in vitro* evaluation of probiotic treatments for nasal *Staphylococcus aureus* decolonization

**DOI:** 10.1101/2022.08.28.505587

**Authors:** Burcu Tepekule, Weronika Barcik, Willy I. Staiger, Judith Bergadà-Pijuan, Thomas Scheier, Laura Brülisauer, Alex Hall, Huldrych F. Günthard, Markus Hilty, Roger D. Kouyos, Silvio D. Brugger

**Author notes:** These authors contributed equally (Markus Hilty, Roger D. Kouyos, Silvio D. Brugger).

## Abstract

**Abstract:** Despite the increasing burden of antibiotic resistance and persistence, current approaches to eradicate nasal pathobionts such as *Staphylococcus aureus* and *Streptococcus pneumoniae* are based on the use of antibacterial agents. An alternative approach is the artificial inoculation of commensal bacteria, i.e., probiotic treatment, which is supported by the increasing evidence for commensal-mediated inhibition of pathogens. To systematically investigate the potential and the limitations of this approach, we developed a quantitative framework simulating the dynamics of the nasal bacterial microbiome by combining mathematical modeling with longitudinal microbiota data. By inferring the microbial interaction parameters using 16S rRNA amplicon sequencing data and simulating the nasal microbial dynamics of patients colonized with *S. aureus*, we compared the decolonization performance of probiotic and antibiotic treatments under different assumptions on patients’ bacterial community composition and susceptibility profile. To further compare the robustness of these treatments, we simulated a *S. aureus* challenge following each treatment and quantified the recolonization probability. Eventually, using nasal swabs of adults colonized with *S. aureus*, we confirmed that after antibiotic treatment, recolonization of *S. aureus* was inhibited in samples treated with a probiotic mixture compared to the non-treated control. Our results suggest that probiotic treatment clearly outperforms antibiotics in terms of decolonization performance, recolonization robustness, and leads to less collateral reduction of the microbiome diversity. Moreover, we find that recolonization robustness is highest in those patients that were not initially colonized by *Dolosigranulum pigrum*. Thus, probiotic treatment may provide a promising alternative to combat antibiotic resistance, with the additional advantage of personalized treatment options via using the patient’s own metagenomic data to tailor the intervention. The combination of an *in silico* framework with *in vitro* confirmatory experiments using clinical samples reported in this work is an important step forward to further investigate this alternative in clinical trials.

**Importance:** The development of new antimicrobial agents is declining while antibiotic resistance is rising, which is particularly concerning for upper respiratory tract pathogens *S. pneumoniae* and *S. aureus*. Combating such resistant infections will only become more challenging unless alternative treatment strategies are explored. Despite the accumulating evidence on using commensal bacteria for pathobiont decolonization, it is still not commonly practiced. To investigate the potential of commensal-mediated inhibition of pathogens systematically, we developed a quantitative framework describing the dynamics of the nasal microbiome by merging mathematical modeling and metagenomic data. We show that probiotic treatment outperforms antibiotics regarding decolonization performance and recolonization robustness while preserving the microbiome diversity with the additional advantage of personalized treatment options via using the patient’s own microbiota data. Moreover, we validated the approach by using nasal swabs from adults with nasal *S. aureus* colonization, demonstrating that probiotic treatment prevents recolonization with *S. aureus in vitro*. The framework developed in this work is an important step forward for the translation of experimental and clinical data into mainstream clinical practice in a systematic and controlled manner.

## 1. Introduction

The global emergence and spread of antimicrobial resistance (AMR) and persistence is of great concern to public health and an important driver of treatment failure, increased morbidity and mortality, and healthcare costs (1). Invasive infection with pathobionts, such as methicillin-resistant *Staphylococcus aureus* (MRSA) and multidrug resistant (MDR) *Streptococcus pneumoniae*, is subsequent to colonization and nasal carriage is a risk factor for several infective syndromes. Despite the increasing problem of AMR and antimicrobial persistence, treatment options are scarce, and current interventions have several limitations (2). In particular, complicated decolonization protocols using antimicrobial substances have low success rates, and promote the emergence of antimicrobial resistance. Therefore, new solutions to control the colonization and thus the infection of drug-resistant and persistent pathogens are urgently needed. One promising alternative is the use of microbiota-mediated therapeutics for decolonization of such resistant and persistent bacteria (3, 4).

In terms of AMR, one of the major threats is the spread of the resistant Gram-positive bacteria such as MRSA and MDR *Streptococcus pneumoniae*, which are associated with poor clinical outcomes (5, 6). The first step to invasive disease caused by the pathobionts *S. aureus* and *S. pneumoniae* is colonization (7). Nasal carriage of these bacteria is a well-documented risk factor for skin and soft tissue infections, pneumonia, surgical-site-infections, and bacteremia and causes significant morbidity and mortality worldwide (8–15). Despite the increasing problem of AMR, current methods to eradicate these pathobionts from the nasal cavity are based on the application of mupirocin for *S. aureus* (16–18). It has been repeatedly shown that mupirocin treatment can promote the emergence of antibiotic resistance in *S. aureus*, including MRSA (19, 20). As a result, non-antibiotic microbiota-targeted therapy seems to be a promising alternative approach based on the increasing data on commensal-mediated inhibition of pathogens.

There has been accumulating evidence on *Corynebacterium* and *Dolosigranulum* species providing colonization resistance against *S. aureus* and *S. pneumoniae* in the nasal cavity of humans (21–30). However, mechanisms by which these common commensal organisms eliminate *S. aureus* or *S. pneumoniae* colonization are not exactly clear (31). A commonly proposed hypothesis suggests that secondary metabolites appear to govern bacterial competition in the human nose and may become important agents for eradication of opportunistic pathogens from human microbiota (31, 32) including bacteriocins, bacteriocins-like antimicrobial substances, and other bioactive molecules(30, 33, 34). Hence, the clinical use of commensal organisms presents an alternative to antibiotic treatment, as studies have shown successful decolonization of *S. aureus* through the artificial inoculation of *Corynebacterium pseudodiphtheriticum* (35) and *Corynebacterium* strain Co304 (36). However, due to the intricate interactions within the microbiota, such interventions can lead to indirect and unforeseen consequences. Therefore, mathematical modeling of commensal-mediated pathogen inhibition is essential: it enables a systematic and controlled investigation of potential outcomes and their determinants, thereby facilitating the development of personalized microbiota- targeted therapies.

Previous research has successfully identified certain qualitative relationships between the key residents of the nasal microbiota (21–30). However, all this information on the community composition and the pairwise interactions still remain mostly on a descriptive level, focusing on the correlations of presence and absence of the microbial species. A complementary approach is to incorporate this data in a dynamical model where the interactions between microbial species could be inferred and followed temporally. This model would allow for complex interactions and feedback loops to result in outcomes that could not have been directly predicted by observing pairwise microbial interactions in isolation. Consequently, it would provide a comprehensive understanding of the temporal dynamics of the nasal microbiota, which has the potential to be manipulated for positive health outcomes as suggested by descriptive clinical studies.

We develop a quantitative framework to design and simulate a treatment regime by the artificial inoculation of beneficial commensals *D. pigrum* and *C. pseudodiphtheriticum* for *S. aureus* and *S. pneumoniae* decolonization. By bringing the qualitative information and the data together in an integrative framework, we provide a quantitative representation of the microbiome dynamics in the nose. Dynamical modeling of the nasal microbiome allows us to make quantitative predictions regarding the effect of clinical interventions such as different decolonization procedures and antibiotic treatment regimens. Moreover, our framework has the potential to be extended as a personalized-treatment tool for precision probiotics, which can be used by physicians in a clinical setting.

## 2. Results

Development of our computational framework can be summarized in three steps: data curation, model building and parametrization, and the implementation of the treatment regimens followed by a *S. aureus* challenge (Fig. 1) (see Methods). Our mathematical model describes the temporal nasal microbial dynamics of a patient already colonized with *S. aureus*, and our simulations include a 5-day long pretreatment regimen with mupirocin in accordance with current eradication protocols followed by either antibiotic or probiotic treatment, and finally a *S. aureus* challenge by the reintroduction of *S. aureus* to the nasal cavity. Our quantitative analysis is based on the comparison of the antibiotic and probiotic treatments by their decolonization performance and their robustness against *S. aureus* recolonization.

**Figure 1:**
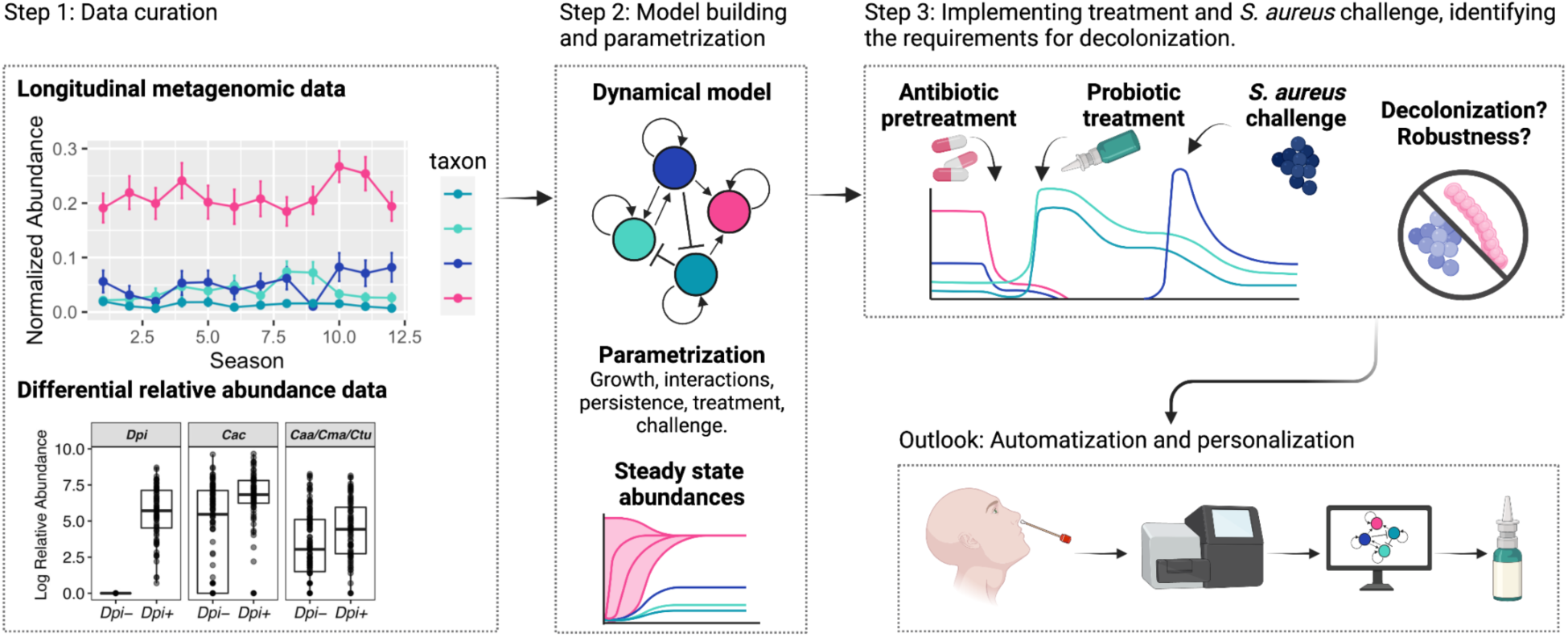
Illustration summarized in three steps: data curation (30, 44), model building and parametrization, and the implementation of the treatment regimens followed by the *S. aureus* challenge. As an outlook, our computational framework can be extended as a personalized-treatment tool for precision probiotics and can be used by physicians in a clinical setting.

After the growth and interaction parameters were inferred, filtered according to the criteria for *S. aureus* and *S. pneumoniae* co-existence, and the relative magnitude of certain inter-oligotypes interaction terms (see Methods), we were left with 954 and 929 parameter sets out of 1000 initially sampled for Population 1 (in which patients are already colonized with *S. aureus* but assumed to harbor no *D. pigrum* prior to treatment) and 2 (in which patients are already co-colonized both with *D. pigrum* and *S. aureus* prior to treatment), respectively. Each of these sets represented a distinct patient and the simulations were repeated for each set for all the combinations of variables which are the number of treatment courses, inoculum of the probiotic cocktail, switching rates between persistent and susceptible phenotypes, and the timing and inoculum of the *S. aureus* challenge. In our model, persister cells represent a subpopulation of cells that can survive antibiotic treatment via going into a dormant state without being resistant (37), i.e., by being temporarily "not susceptible". An example of the dynamics of the mathematical model is presented in Fig. 2, where the same system is simulated with and without persisters for antibiotic and probiotic treatments. These simulations suggest that persisters may represent an obstacle for achieving *S. aureus*

**Figure 2:**
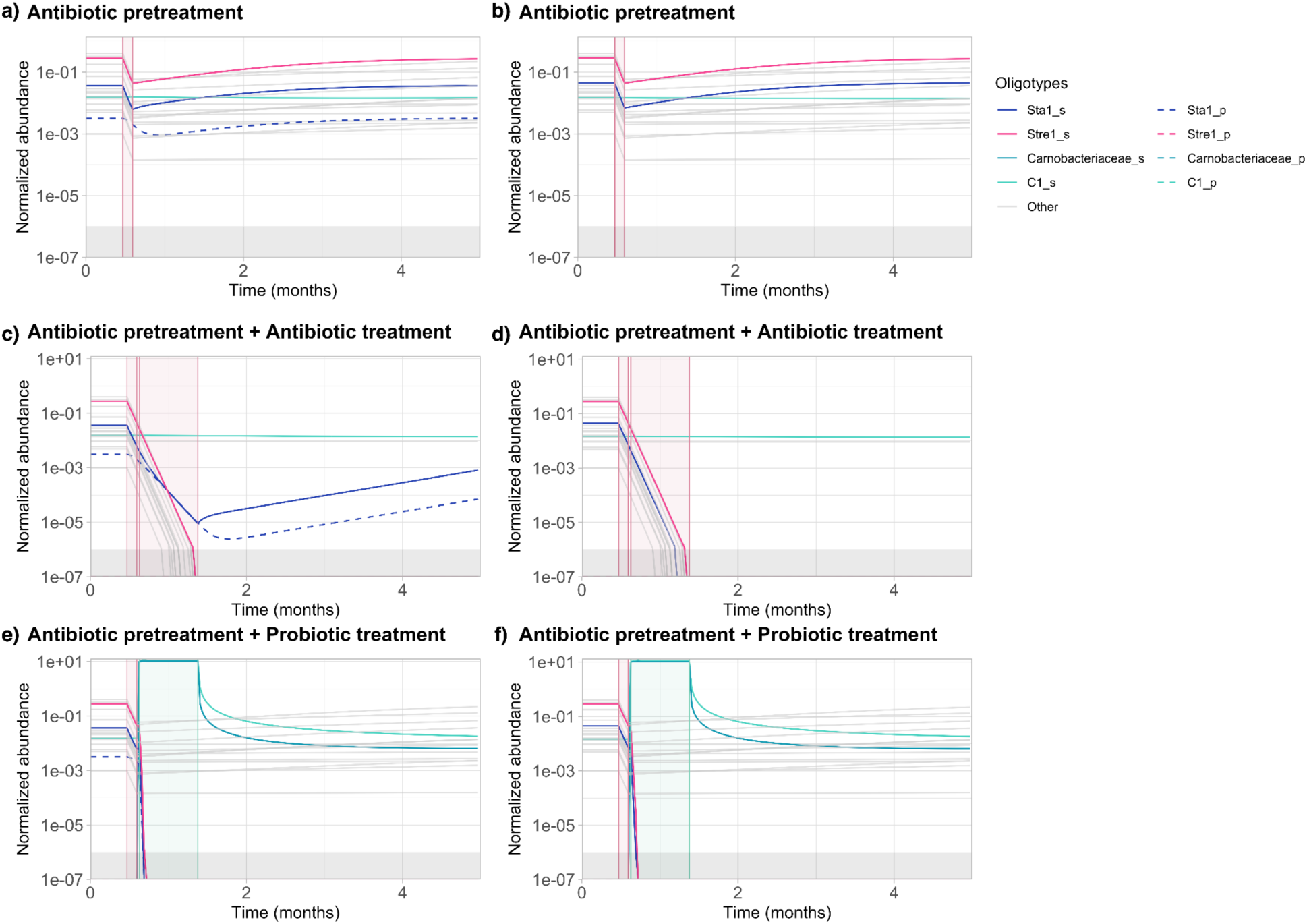
An example of the dynamics of the mathematical model, where the same system is simulated **a)**, **c)**, **e)** with and **b)**, **d)**, **f)** without persisters for only pretreatment, pretreatment followed by antibiotic treatment, and pretreatment followed by probiotic treatment, respectively. s and p suffixes in the figure legend represent the susceptible and persistent counterparts of the same oligotype, respectively. See Table 1 for the IDs used for each oligotype. Gray area below the normalized abundance of 10^−6^ represents the region below the extinction threshold.

decolonization by antibiotic treatment whereas the probiotic treatment is more robust to persisters. We therefore start by systematically considering the more conservative assumption of systems without *S. aureus* persisters and then assess the impact of different persister frequencies.

### Probiotic treatment has a higher probability of *S. aureus* decolonization *in silico*

In the case of no persisters, over all the inoculum sizes and number of treatment courses, probiotic treatment considerably outperforms antibiotic treatment in terms of *S. aureus* decolonization, except for a few cases where the normalized inoculum size of the probiotic treatment is very low (10^−1^) and the number of treatment courses is very high (> 5) (Fig. 3a)). Even in those cases, probiotic treatment leads to a probability of decolonization above 0.9, and the difference in decolonization probability between the antibiotic and the probiotic treatment remains below 0.09. Moreover, a single course of probiotic treatment leads to a decolonization probability between 0.41 to 0.99 depending on the inoculum size whereas a single course of antibiotic treatment never leads to decolonization.

**Figure 3:**
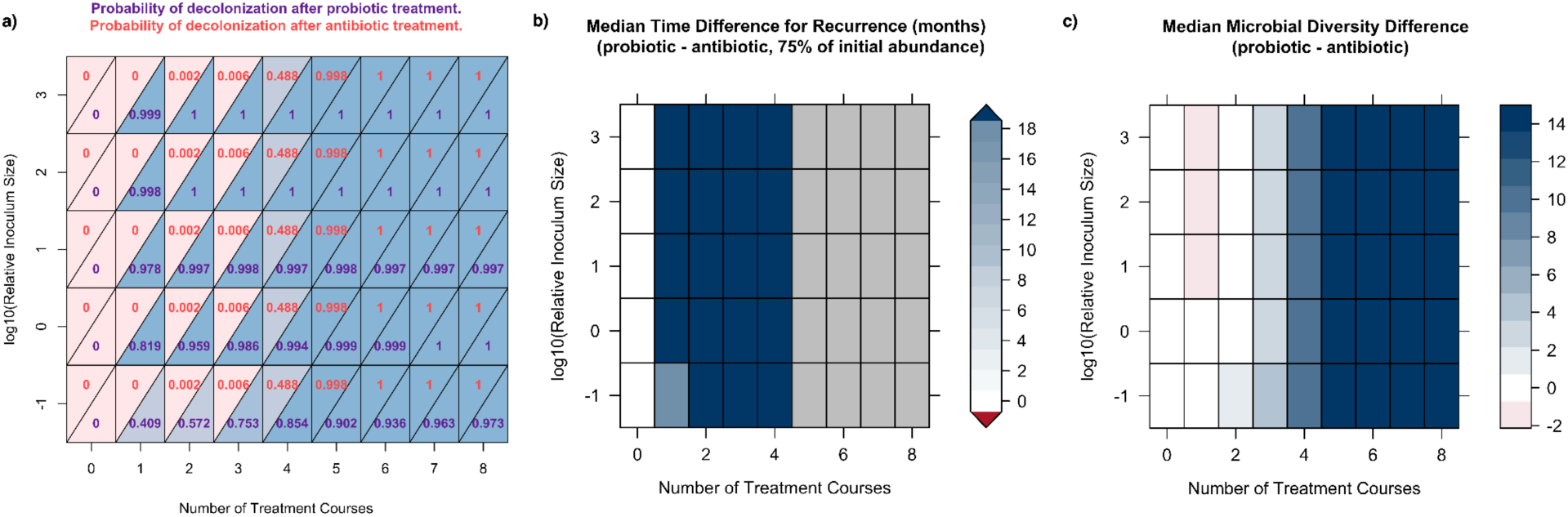
Decolonization probability, time to recurrence, and final microbial diversity results for Population 1, with no persisters. **a)** Probability of *S. aureus* decolonization for different number of treatment courses and normalized inoculum sizes. **b)** Median time difference for recurrence between probiotic and antibiotic treatments. **c)** Median microbial diversity difference between probiotic and antibiotic treatments, calculated as the difference of the number of oligotypes still present in the microbiome following treatment. Gray boxes in **b)** represent the cases where antibiotic and probiotic treatments have a median time recurrence longer than 36 months, and therefore the difference cannot be calculated. See Supplementary Fig. S1 for the same analysis for Population 2.

### Antibiotic treatment leads to faster *S. aureus* regrowth when decolonization fails

To quantify how fast *S. aureus* grows back in case of an unsuccessful decolonization procedure, we checked the time it takes for *S. aureus* to reach 75% of its initial abundance prior to any treatment within the next 36 months following the treatment regimen, which we refer to as *S. aureus* recurrence. Median values of the difference in time to recurrence for the probiotic and the antibiotic treatment are given in Fig 3b). For the probiotic treatment, recurrence within the next 3 years following treatment only occurs for the combination of minimum normalized inoculum size (10^−1^) and the minimum number of treatment courses (1) and takes a median value of 21.9 months (standard deviation (sd) 8.5 months). We found that allowing for *S. aureus* persisters substantially increases the benefits of probiotic treatment compared to antibiotics (Fig. 4). When all combinations of switching rates (including no persisters with a switching rate of 0) are considered, the time to recurrence after probiotic treatment takes a median value of 29.5 months (sd=8.3 months) (Fig. 4a)), assuming that persister *S. aureus* cells are equally affected as the susceptible *S. aureus* cells by the inoculum of *D. pigrum* and *C. pseudodiphtheriticum* (see Methods). On the contrary, recurrence within the next 3 years following antibiotic treatment is observed up to 4 treatment courses (20 days of treatment) for all switching rates including 0 (case of no persisters) and for higher switching rates, recurrence can occur up to 7 treatment courses (35 days of treatment) (Fig. 4b)).

**Figure 4:**
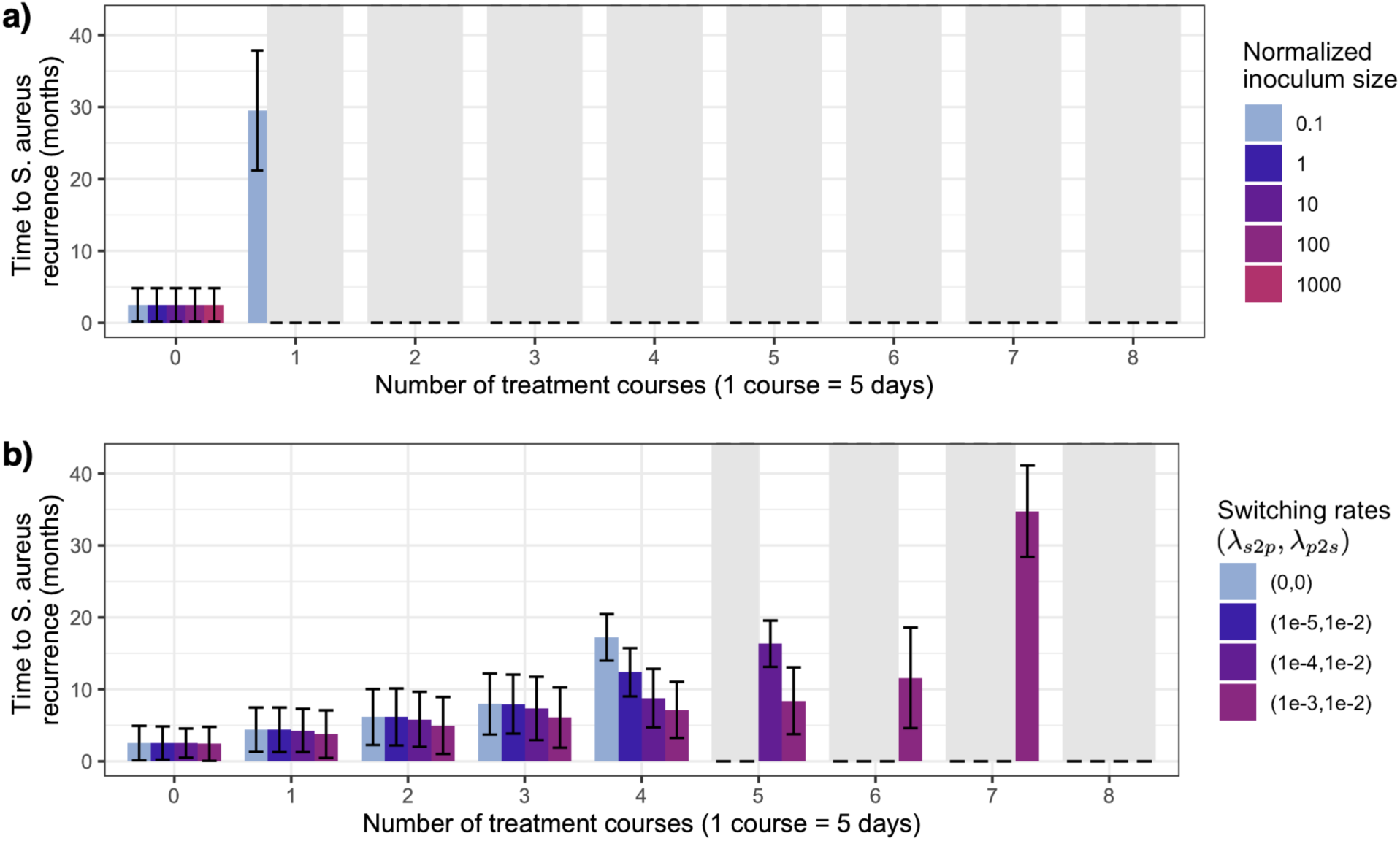
Time to *S. aureus* recurrence for Population 1, with different normalized inoculum sizes for the probiotic (**a)**), and different levels of persisters for the antibiotic (**b)**) treatments. Gray bars represent durations with a mean value longer than 36 months, meaning that no recurrence is observed within the 3 years following treatment. Error bars display +/- the standard error on the mean recurrence time. Note that unlike the probiotic treatment, the antibiotic treatment does not utilize inoculation of commensals, therefore the analysis is based on varying levels of persisters only. Similarly, persisters are assumed to play no role during probiotic treatment, thus the analysis is based on varying levels of inoculum size only. See Supplementary Fig. S2 for the same analysis for Population 2.

### Antibiotic treatment substantially reduces nasal microbiome diversity at the cost of *S. aureus* decolonization

Antibiotic and probiotic treatment leads to equal probabilities of decolonization when the number of treatment courses is high enough (> 5 courses for either antibiotic or probiotic treatment) in the case of no persisters (Figs. 3a) and b)). However, longer durations (i.e., higher number of treatment courses) of antibiotic treatment affect the other members of the nasal microbiome as well, reducing the number of oligotypes present prior to the treatment. In the case of no persisters and within the region where both antibiotic and probiotic treatment leads to equally successful decolonization, the difference between the number of oligotypes still present in the nasal microbiome following treatment has a median value of 14 (Fig. 4c)), with probiotic treatment having 16 and antibiotic treatment having 2 oligotypes left on average. Note that this calculation also includes *S. aureus* and *S. pneumoniae* as oligotypes, thus leading to a median difference of -2 when the number of treatment courses is equal to 1 and where probiotic treatment still leads to a successful decolonization of *S. aureus* and *S. pneumoniae* for higher inoculum sizes whereas antibiotic treatment fails to do so.

### Probiotic treatment is more robust to subsequent challenge with *S. aureus* regardless of *S. pneumoniae* extinction

An example of the dynamics of the *S. aureus* challenge is presented in Fig. 5. Figs. 5a) and c) represent a patient where the challenge does not lead to *S. aureus* regrowth following the probiotic treatment (hence leads to extinction) but leads to *S. aureus* recolonization following the antibiotic treatment. Figs. 5 b) and d) represent a patient where the challenge leads to *S. aureus* recolonization following both the antibiotic and the probiotic treatments. The probability of *S. aureus* extinction following the *S. aureus* challenge is used as a metric to assess the robustness of each treatment. For population 1 with no persisters, probiotic treatment had an average extinction probability of 46.1% and was exceedingly more robust to *S. aureus* challenge than antibiotic treatment which had an average extinction probability of 0.2% (Fig. 6a)), where the extinction probabilities are averaged across the number of treatment courses, normalized inoculum size of the probiotic cocktail, and the normalized inoculum size and the time of the *S. aureus* challenge. On the contrary, for population 2 with no persisters, probiotic treatment had an average extinction probability of 2.2% and the antibiotic treatment had an average extinction probability of 1.3% (Fig. 6c)). We segregated the robustness of each treatment by *S. pneumoniae* extinction for each population since *S. pneumoniae* is one of the major competitors of S. aureus and also targeted by both the antibiotic and probiotic treatments. For the antibiotic treatment, given that *S. pneumoniae* goes extinct during treatment, the probability of S*. aureus* extinction in the case of a *S. aureus* challenge decreases substantially for both populations (Figs. 6a) and b) for population 1, Figs. 6c) and d) for population 2). For population 1, the robustness of the probiotic treatment is affected by *S. pneumoniae* extinction less than antibiotic treatment does (Figs. 6a) and b)) and is almost indifferent to *S. pneumoniae* extinction for population 2 (Figs. 6c) and d)).

**Figure 5:**
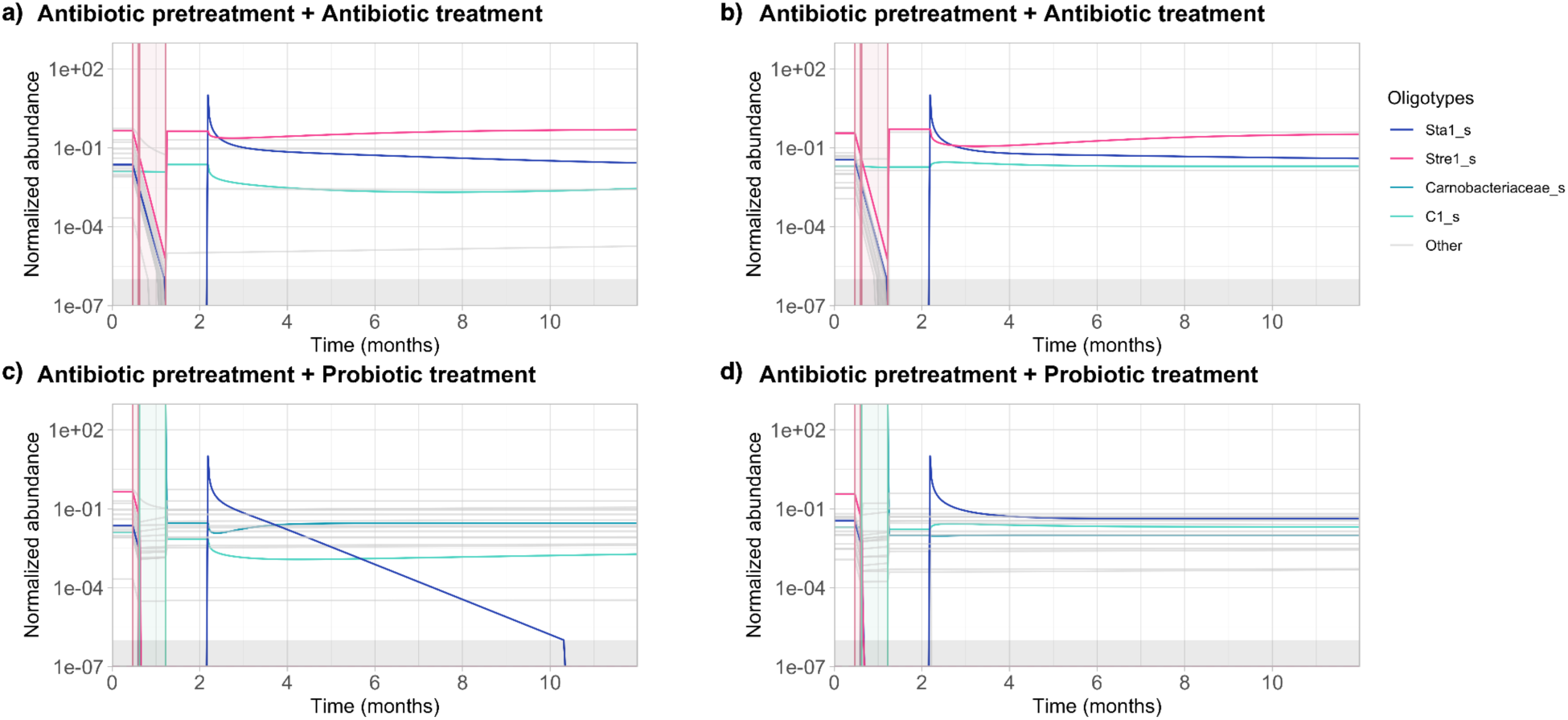
An example of the dynamics of *S. aureus* challenge for Population 1, where **a)** and **c)** and **b)** and **d)** belong to two different patients. *S. aureus* challenge leads to recolonization for both patients following antibiotic therapy (**a)** and **b)**), and for the second patient following probiotic therapy (**d)**), whereas *S. aureus* fails to regrow following probiotic treatment for the first patient (**c)**). Gray area below the normalized abundance of 10^−6^ represents the region below the extinction threshold.

**Figure 6:**
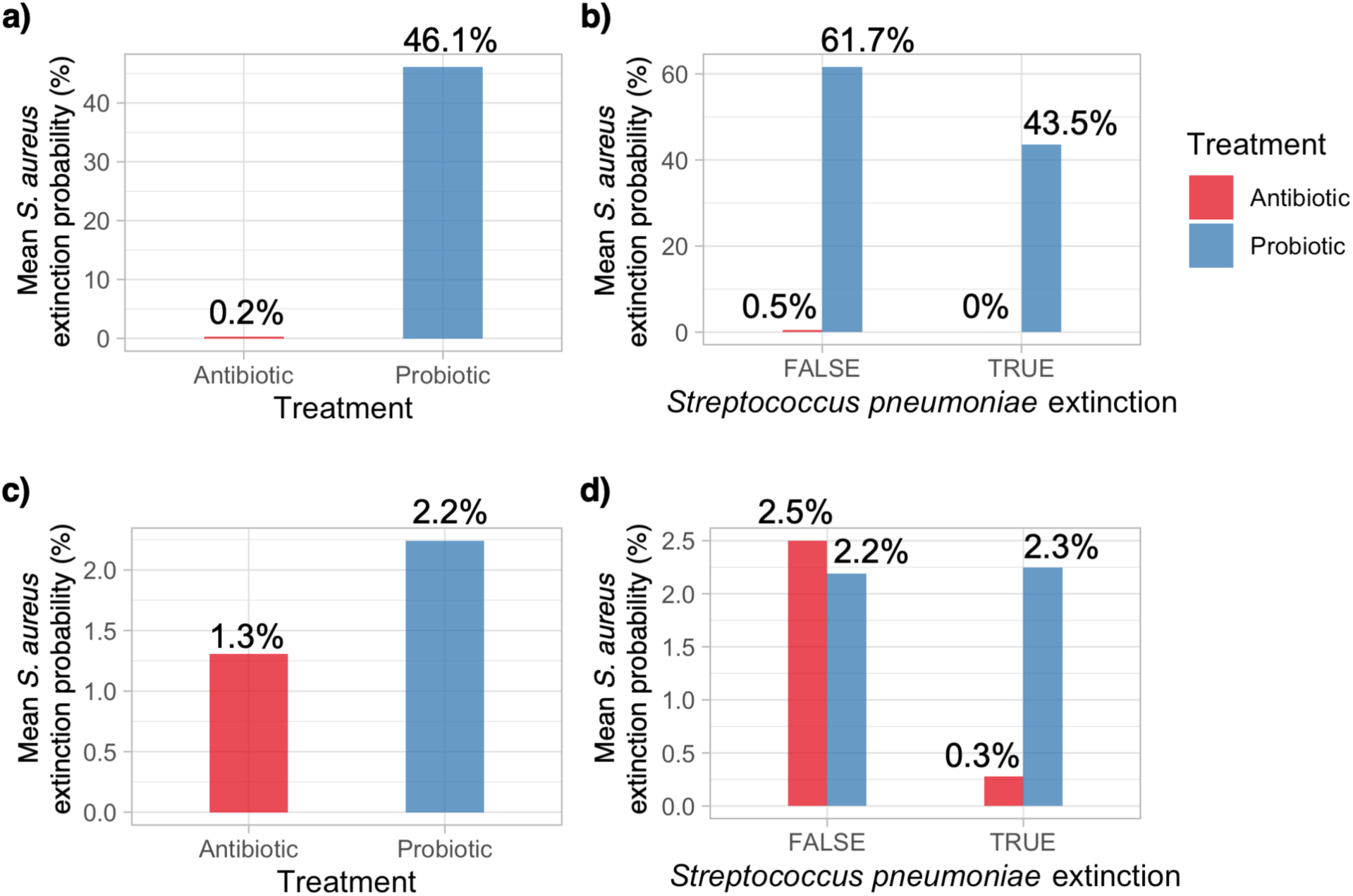
Mean *S. aureus* extinction probabilities for the antibiotic and probiotic treatments followed the *S. aureus* challenge **a)**, **b)** for Population 1, and **c)**, **d)** for Population 2; **b)**, **d)** where the probabilities are segregated by *S. pneumoniae* extinction.

### Probiotic treatment prevents recolonization of nasal microbiota with *S. aureus in vitro*

To test the framework, we designed an experimental *in vitro* setup to evaluate commensal probiotic treatment to prevent *S. aureus* recolonization. Nasal swabs were collected from seven methicillin- susceptible *S. aureus* (MSSA) carriers. These swabs were then treated with mupirocin to mimic patient nasal decolonization. Afterwards, the swab fluid was divided into two aliquots. One aliquot was treated with a probiotic cocktail containing *D.pigrum* and *C. pseudodiphtheriticum* in a 1:1 ratio, while the other served as the non-treated control. To assess *S. aureus* recolonization probability, aliquots of MSSA, treated either with probiotics or left untreated, were challenged with MRSA (Fig. 7 a)). We found that recolonization was significantly lower (reaching up to a 5 log10 reduction in CFU/ml) in aliquots treated with the probiotic cocktail compared to non-treated aliquots (Fig. 7 b)). This behavior was observed in all donors (n=7). In one donor, however, the magnitude of the effect was significantly lower (2.44 times lower recolonization number (CFU/ml) in the probiotic treated aliquot compared to the control). The experiment for this specific donor was repeated and showed a consistent reduction in the probiotic treated aliquot, however still lower than the average (Fig. 7 b)). These results exceed the predictions of the mathematical model of an estimated 46.1% extinction rate of *S. aureus* upon challenge (Fig. 6 a)).

**Figure 7:**
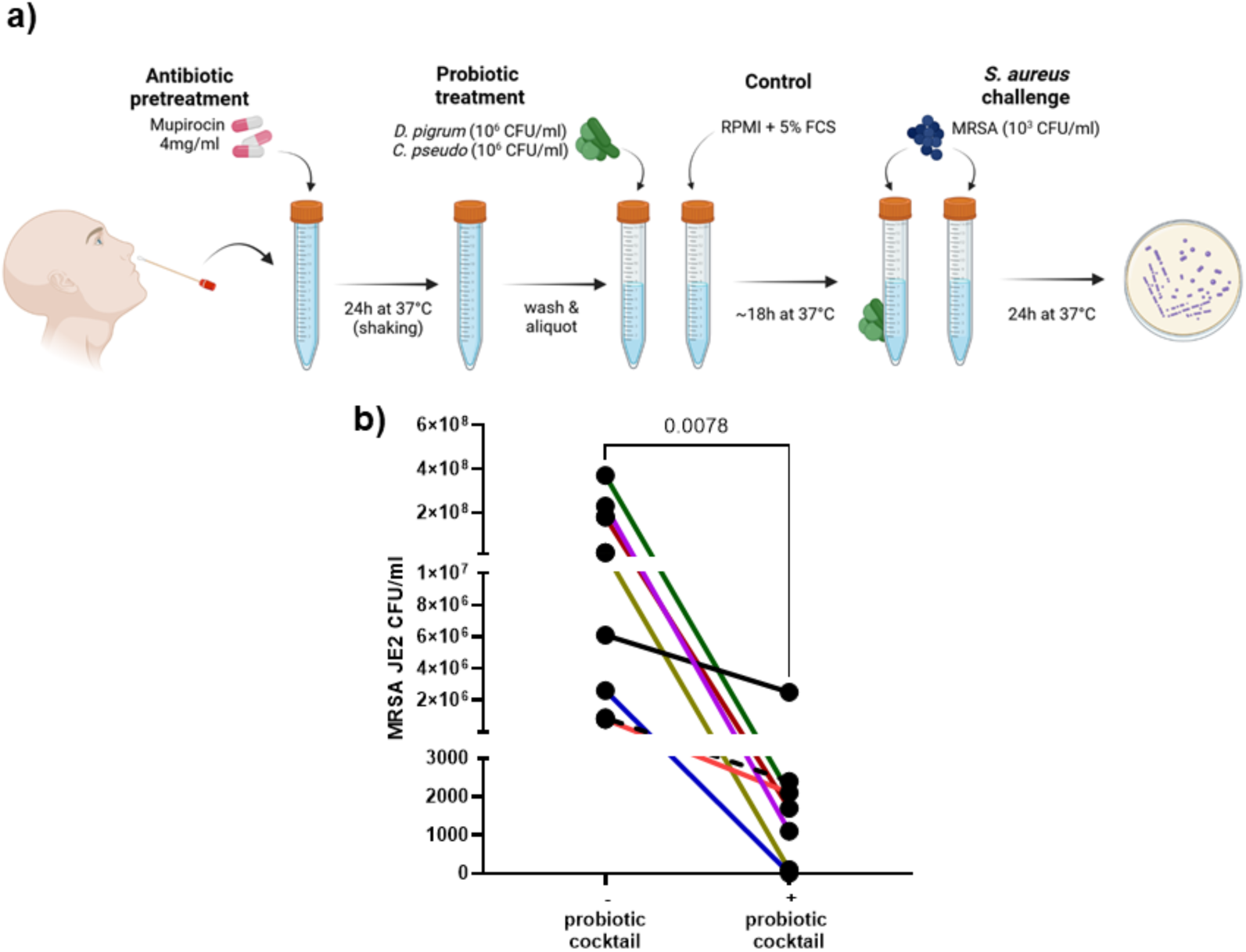
Recolonization of MRSA in CFU/ml in the nasal swab liquid aliquots treated with probiotic cocktail containing *D. pigrum* and *C. pseudodiphtheriticum* in a 1:1 ratio versus the untreated aliquots. Scheme of the *in vitro* experiments (**a)**) that show significantly lower recolonization of MRSA in aliquots supplemented with the probiotic cocktail (**b)**) compared to untreated aliquots. Note that for one donor, two replicates (represented by the black line and black dashed line) were included in the analysis (see main text). See supplementary figures for individual replicate analysis (Fig. S5), that remained the same as for the pooled analysis. For statistical analysis Wilcoxon matched-pairs signed rank test was used. Scheme of the experiment was generated with BioRender.

## 3. Discussion

Despite accumulating evidence supporting the use of commensal bacteria for pathobiont decolonization, it remains an underutilized alternative to antibiotic treatment in combating resistance and persistence. To investigate the potential of this alternative approach based on commensal- mediated inhibition of pathogens in a systematic way, we developed a quantitative framework describing the dynamics of the nasal microbiome by merging mathematical modeling and the data provided in the literature.

Our results suggest that on average, probiotic treatment has a higher chance of sustainable *S. aureus* decolonization when compared to antibiotic treatment, even for *D. pigrum* and *C. pseudodiphtheriticum* inoculum sizes lower than commonly practiced in a clinical setup (Fig. 3a)). Moreover, in an experimental *in vitro* system, we investigated the level of recolonization of *S. aureus* after probiotic treatment and we observed significant inhibition of pathobiont (Fig. 7b)), supporting findings from our mathematical modeling approach.

When persister *S. aureus* cells are included in the model, antibiotic therapy leads to an even lower chance of decolonization and shorter time to *S. aureus* recolonization when the decolonization is unsuccessful (Fig. 4b)) while probiotic therapy is robustly effective. The choice of adding persister cells to our model stems from the frequently observed phenotypic heterogeneity among the patient- derived *S. aureus* strains (38). Since the mechanisms of how *D. pigrum* and *C. pseudodiphtheriticum* inhibit the growth of *S. aureus* are not exactly clear (39), we assumed that persister *S. aureus* cells are equally affected as the susceptible *S. aureus* cells by the inoculum of *D. pigrum* and *C. pseudodiphtheriticum* (40). As a result, introduction of persistence to the model only lowers the average growth rate of *S. aureus* for the probiotic treatment, thus not having a substantial effect on its decolonization performance. Although this assumption favors probiotic treatment in terms of decolonization performance, it would not affect its robustness against *S. aureus* challenge since for most of the parameter sets, the new microbial community composition after probiotic treatment does not allow for *S. aureus* to have a positive growth rate (Fig. 6a) and b)).

Comparing the antibiotic and the probiotic treatment purely by their decolonization performance masks the collateral damage induced by antibiotics when both treatments perform equally well. In fact, when the number of treatment courses is high enough, i.e., the treatment is continued for a sufficient amount of time, both treatments lead to decolonization almost with certainty (Fig 3a)). However, such an aggressive application of antibiotics drives other beneficial members of the nasal microbiome to extinction, thus reducing the overall microbial diversity in the nasal cavity (Fig. 3c)). Loss in microbial diversity makes the host more susceptible to the effects of environmental exposures and leads to an increased risk of colonization by pathogenic species including recolonization by *S. aureus* itself (41). Loss of microbial diversity is substantially lower for probiotic treatment even for a high number of treatment courses, allowing the host to still maintain the beneficial bacteria in their nasal cavity which is essential for upper respiratory tract health.

Due to the impact both treatments have on the microbial community composition, we also investigated how the host would respond to a *S. aureus* challenge after the treatment has ended. Our results suggest that probiotic treatment is more robust to *S. aureus* challenge, i.e., considerably more successful in preventing *S. aureus* recolonization (Fig. 6a)), providing the host with recolonization resistance for future *S. aureus* infections. Even for cases where probiotic treatment also decolonized *S. pneumoniae*, which is one of the major competitors of *S. aureus*, the robustness of the probiotic treatment was maintained (Fig. 6b)). On the contrary, antibiotic treatment barely provided any recolonization resistance, supporting the argument for its short term protection and negative collateral effects on microbial diversity and robustness.

Results above were obtained for Population 1, where *D. pigrum* and *S. aureus* co-existence is not imposed on the model based on the inverse association between *S. aureus* carriage and the presence of *D. pigrum* in the nasal microbiome (21–26, 42). This means that the model is indifferent to *D. pigrum* and *S. aureus* co-existence, and the patients do not harbor *D. pigrum* in their nasal microbiome prior to any treatment. To test the robustness of the probiotic treatment even further, we repeated our *S. aureus* challenge analysis on Population 2, which has the constraint of *S. aureus* and *D. pigrum* co-existence in the nasal microbiome prior to treatment. Although less realistic, it is a more difficult patient population to treat from the probiotic treatment’s perspective since the model parameters allow some amount of the treatment itself, i.e., *D. pigrum*, to already co-exist with the pathogen, i.e., *S. aureus*, prior to the application of probiotics. Unsurprisingly, robustness of the probiotic treatment was substantially lower for Population 2 but was still higher compared to the antibiotic treatment (Fig. 6c)). Segregation of the results by *S. pneumoniae* decolonization led to similar results as with Population 1 and revealed that the low levels of colonization resistance provided by the antibiotic treatment can mostly be attributed to *S. pneumoniae* still being a part of the nasal microbiome (Fig. 6d)), which is also a risk factor for pneumonia, otitis media, sepsis, etc.

Although having two different patient populations provided valuable insights on treatment robustness, inferring interaction parameters for *D. pigrum* using a sample taken from a patient belonging to Population 1 is theoretically impossible. Since *D. pigrum* is assumed to be absent prior to any treatment, its interaction with other oligotypes, including itself in the nasal microbiome, cannot be estimated, imposing an epistemic limitation on parameter inference. We circumvented this limitation via sampling from posterior distributions obtained by using samples where *D. pigrum* is present in the nasal cavity prior to treatment, implicitly assuming that interaction parameters involving *D. pigrum* would be similar when *D. pigrum* and *S. aureus* can co-colonize the nasal cavity to a certain extent.

Aligning with the mathematical model, our *in vitro* experimental data show an increased robustness against *S. aureus* (MRSA) challenge after antibiotic pretreatment followed by probiotic treatment compared to antibiotic treatment alone. Furthermore, the strong reduction of *S. aureus* growth (in 6 out of 7 donors, Fig. 7b)) exceeded the predictions of 46.1% given by our mathematical model (Fig. 6a)). In one donor, however, only a slight decrease was detected. We suspect that several factors might have influence on the level of *S. aureus* reduction: one of them could be initial composition of the microbiota before antibiotic pretreatment and, as a consequence, composition and abundance of the remaining microbiota after antibiotic pretreatment. The example of the latter was shown in our mathematical model (Fig 6b)) where an influence on the robustness against *S. aureus* challenge was shown for *S. pneumonia*. Nasal microbiota compositions differ not only between individuals (43) but also in the individual itself over time (22), thus explaining the variability seen after the repeat of the one donor.

The 16S rRNA gene sequencing data used in our study was based on the most detailed longitudinal dataset we had access to (provided in (44)), allowing us to build a more detailed model regarding temporal microbial community dynamics. However, this data was based on the samples taken from infants, whereas our probiotic treatment regimen is targeted at adults. This represents, however, only a minor limitation for our study, as this data is used only to estimate the interaction parameters between the microbial species and the steady state of the system using varying initial conditions, but not the longitudinal relative abundances themselves. Given that the negative interactions between the commensals (*D. pigrum* and *C. pseudodiphtheriticum*) and the pathobionts are reported in the same direction in adults (29, 45–47) and children (25, 26, 48–53), we believe it would be plausible to assume that these bilateral interaction parameters are similar between adults and infants older than 12 weeks of life who are not significantly impacted by the maternal nasal microbiome.

Despite the indisputable advantages of probiotic treatment suggested by our results, our analysis is conservative in terms of favoring probiotic treatment. Our model imposes an equal inoculum and identical timing on *D. pigrum* and *C. pseudodiphtheriticum* application, which can easily be modified and has the potential to improve the decolonization outcomes for shorter treatment durations. Furthermore, such modifications can be based on the metagenomic data of the patient and can even be personalized. Our analysis, including two patient populations depending on their *D. pigrum* colonization prior to treatment, already provides important insights on how sensitive the nasal microbiome might be to recolonization by an environmental *S. aureus* exposure, thus informing the clinicians to take further prevention measures depending on the specific patient.

Although some concerns regarding the safety of probiotics targeting the gut microbiome were raised by several clinical trials (54), studies based on the nasal application of probiotics only revealed minor side effects (55) and are considered to be safe (30, 56–58). On the contrary, the application of antibiotics is quite detrimental to the overall microbial diversity (59), contributing to the emergence and spread of antibiotic resistance and thus jeopardizing the success of the future treatments that would be necessary (60).

Given the public-health crisis associated especially with MRSA, it is obvious that we have to accelerate the process of developing alternative treatments. This is only possible if modeling goes hand in hand with clinical practice, establishing an iterative flow of information. The quantitative framework on commensal-mediated inhibition of pathogens provided in this study is a crucial step forward for the translation of experimental and clinical data into mainstream clinical practice in a systematic and controlled manner. Eventually, validation of the proposed model either *in vivo* or *in situ* is essential for verifying its robustness and practical applicability. However, by laying out the qualitative predictions before any validation via clinical trials, we decouple the model structure from data, which permits us to truly challenge the mechanistic assumptions of the model, which could not be possible if the model was informed by clinical data prior to validation. The theoretical approach proposed in this work could be used both for guiding strategies to treat disease and to inform clinical trial design to optimize human microbiota composition to prevent pathogen colonization and thus to prevent bacterial disease. All these clinically relevant outcomes would also lower the economic burden of antibiotic treatment and healthcare costs and provide the option of a personalized treatment instead of a “one-fits-all” approach.

## 4. Methods

Our computational framework simulates the nasal microbial community dynamics of a patient who is already colonized with *S. aureus*, and goes through either probiotic or antibiotic treatment for decolonization. As the conventional clinical practice, a 5-day long pretreatment regimen with mupirocin is prescribed prior to both treatments. After the pretreatment, the patient either continues to use mupirocin (antibiotic treatment) or applies a probiotic cocktail of *D. pigrum* and *C. pseudodiphtheriticum* to the nasal cavity on a daily basis for *S. aureus* decolonization (probiotic treatment). The patient is challenged with a certain inoculum of *S. aureus* after the treatment is complete to further assess the robustness of the respective treatment to recolonization. Considering the role of antibiotic-persistent bacteria in severe invasive infections, which are often difficult to treat, all simulations are repeated with different levels of antibiotic-persistent *S. aureus* to understand the role of persistence in pathogen decolonization. Development of our framework is summarized in Fig. 1.

### 4.1. Data Curation

To build and parameterize our dynamical model, we used the 16S rRNA gene sequencing data obtained from the nasal swabs of healthy infants (provided in (44)). This data contained relative abundances of nasal microbial oligotypes belonging to 47 infants in total, which had to be filtered and aligned in time before it could be used for parameterization. First, data belonging to the first 12 weeks of life is discarded to avoid any bias that can be introduced by the maternal nasal microbiome. Afterwards, we identified the infants with a nasal microbiome composition such that *S. aureus* and *S. pneumoniae* constitute at least 25% on average of the overall *Staphylococcaceae* and *Streptococcaceae* abundance within the time frame the samples were taken to represent the nasal biota of the already colonized individuals. This abundance threshold is determined heuristically to avoid numerical issues in the inference procedure related to extinction events. After the selection of the infants was complete (11 subjects were left out of 47), their samples were aligned in time according to the month they were taken to represent the seasonal aspects of the microbial abundance distribution. Key table for the oligotypes included during model building is given in Table 1 as described in Mika *et al.* (44, 61). Assuming that the biomass of the samples were constant, relative abundance values are considered to represent the normalized abundance values normalized by the total biomass of the nasal cavity.

**Table 1.**
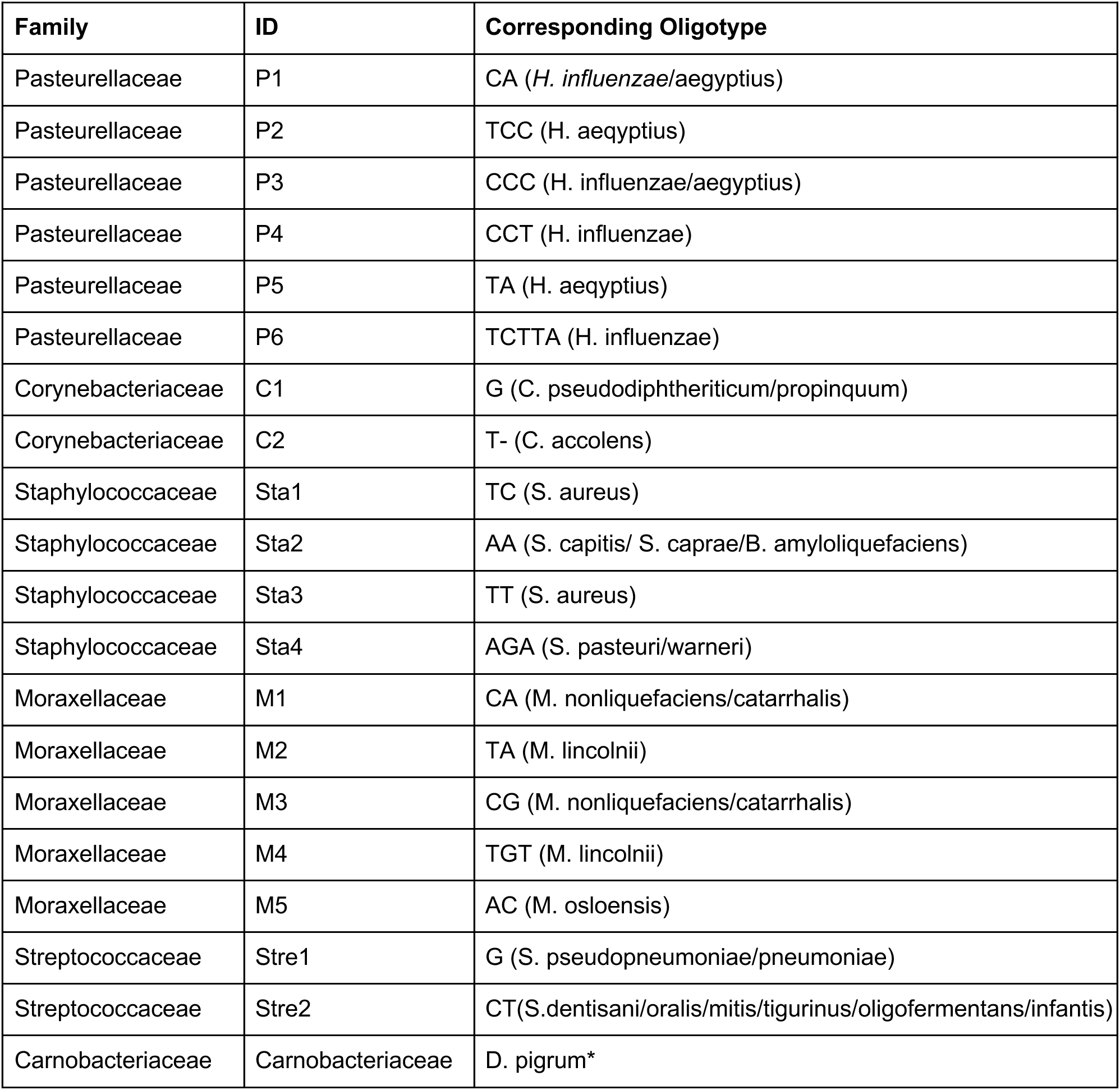
Key Table for Oligotypes belonging to the longitudinal 16S rRNA gene sequencing data. *Since oligotyping was not performed for *D. pigrum*, total *Carnobacteriaceae* abundance is assumed to represent *D. pigrum* abundance, as it is the only member of the family.

Simultaneously, we have integrated the following qualitative data sources to obtain a priori information on the directionality and the relative magnitude of the interactions between our focal nasal microbial oligotypes:

1. ***C. pseudodiphtheriticum* promotes the growth of *D. pigrum* (positive interaction term).** Based on the experimental results reported by Brugger *et al.*, *D. pigrum* benefits from the presence of *C. pseudodiphtheriticum* in terms of growth, but *C. pseudodiphtheriticum* is mainly unaffected by the presence of *D. pigrum in vitro* (30). This suggests a relatively mono-directional interaction between these two commensal species.
2. ***S. pneumoniae* and *S. aureus* inhibit the growth of each other (negative interaction term).** The negative association between *S. aureus* and *S. pneumoniae* is well documented (62, 63) and hypothesized to be due to H^2^O^2^ (64, 65) and/or the pilus produced by *S. pneumoniae in vitro* (66). On the other hand, data from animal models show that this negative association cannot be explained by H^2^O^2^-mediated killing of co-occurring bacteria (67). Consequently, despite the evidence for this negative association, the underlying mechanism remains unclear (39). Therefore, this inhibition is represented in the interaction terms rather than any other additional mechanism.
3. ***D. pigrum* inhibits the growth of *S. pneumoniae* and *S. aureus* (negative interaction term), *C. pseudodiphtheriticum* inhibits the growth of *S. pneumoniae* and *S. aureus* (negative interaction term), and *D. pigrum* inhibits the growth of *S. aureus* more than it inhibits the growth of *S. pneumoniae*.**

There has been accumulating evidence on *Corynebacterium* and *Dolosigranulum* species providing colonization resistance against *S. aureus* and *S. pneumoniae* in the nasal cavity of humans. Multiple studies reported a higher relative abundance of *Corynebacterium* species in people who are not colonized with *S. aureus* (21–24). It has also been repeatedly demonstrated that *Corynebacterium* and *Dolosigranulum* spp. abundances are negatively associated with *S. pneumoniae* colonization in infants (25, 26). Airway microbiota dominated by *Corynebacterium* plus *Dolosigranulum* genera are associated with favorable clinical outcomes compared to microbiota dominated by more pathogenic bacteria such as *Staphylococcus* and *Streptococcus* (27). Similarly, *Corynebacterium* and *Dolosigranulum* were negatively associated with *S. pneumoniae* carriers compared to non-carriers (28). It has been shown that the absolute abundance of *Dolosigranulum* species is a predictor of *S. aureus* carriage in a threshold­dependent manner (29) and *D. pigrum* alone inhibits *S. aureus* but *D. pigrum* plus *C. pseudodiphtheriticum* together inhibits *S. pneumoniae in vitro* (30).

Apart from the assumptions above, we have not imposed any constraints on either the directionality or the magnitude of the remaining interaction terms. Note that these assumptions are qualitative and do not provide any information regarding the quantitative value of the growth and interaction terms. To infer those values, i.e., the rate of growth of taxon !, and the numerical impact of the abundance of one taxon i on the abundance of another taxon ", we have utilized the longitudinal data obtained from the nasal swabs as described in the following sections.

### 4.2. Model building and Parametrization

**The dynamical model.** We used a generalized Lotka-Volterra (gLV) model with a total of 40 compartments representing the normalized abundance of the oligotypes in Table 1, including their susceptible and persister counterparts. Our model is given by the Eqs. 1–2 and the model parameters and variables with their corresponding descriptions are provided in Table 2. Total microbial load in the nasal cavity is assumed to be 10^6^ colony-forming units (CFU), meaning that the extinction threshold for normalized abundances is set to 10^−6^.

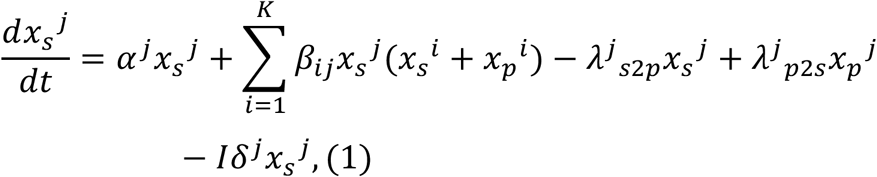

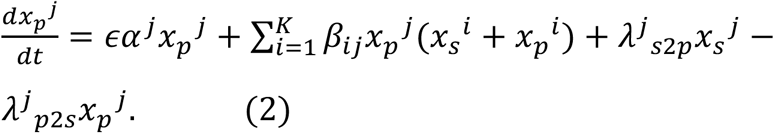

Note that although our analysis focuses mainly on the abundance of the focal species, which are the probiotic commensals and the pathobionts, other oligotypes given in Table 1 are indispensable from an ecological perspective due to the emergent dynamics of the system mediated through the inter-oligotypes interactions.

**Table 2.**
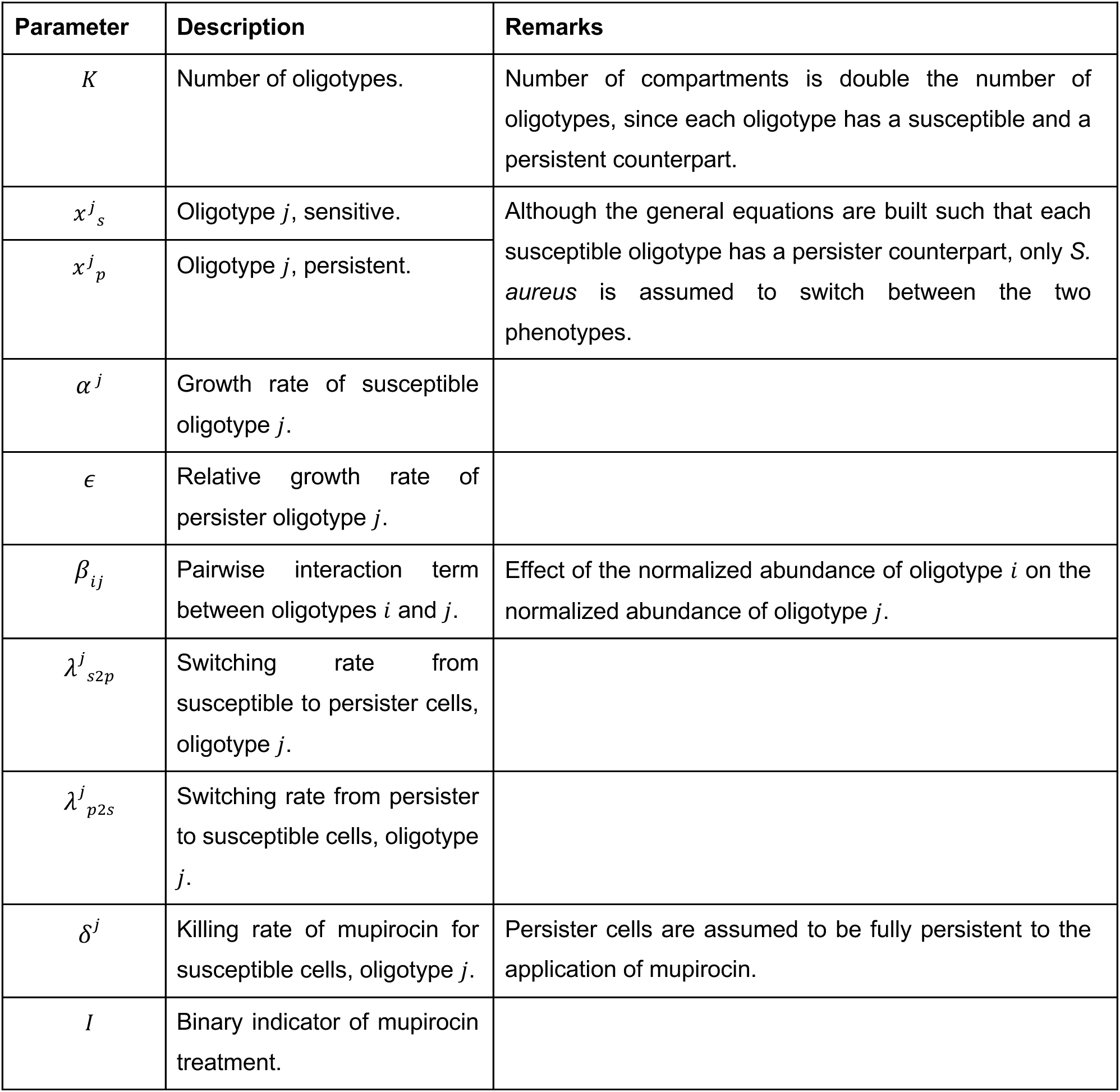
Descriptions of the parameters and variables used in the mathematical model given by Eqns. 1–2.

Growth and interaction parameters. To estimate the growth rates and interaction parameters using the longitudinal 16S rRNA gene sequencing data, we adapted the MDSINE package provided in (68), which is a Markov Chain Monte Carlo (MCMC) based algorithm. Although this estimation algorithm restricts growth rates to have positive and intra-oligotypes interaction terms to have negative values, there are no constraints on other interaction terms. To impose our assumptions provided in the previous section, we manipulated the algorithm so that certain interaction terms are also restricted in their directionality. Instead of using the mean values of the posterior distributions of the growth and interaction terms, we sampled these parameters from the posterior distributions to represent the variability in the parameter space (Detailed information regarding the estimated parameters and convergence diagnostics are provided in Supplementary Table S1 and Supplementary Figs. S3 and S4, respectively). By doing so, we produced a population of models with different quantitative values for each parameter, but with identical constraints. This is analogous to a patient population with the same eligibility criteria for a clinical trial, i.e., a set of simulations using the same dynamical model but different interaction matrices and growth rates sampled from the same posterior distributions.

### Treatment parameters

Killing rate of mupirocin is adjusted such that on average 99% of all the oligotypes except the ones in the family *Corynebacteriaceae* (C1 and C2) are cleared during pretreatment (69, 70). Effect of *D. pigrum* and *C. pseudodiphtheriticum* on *S. aureus* are determined by the inter-oligotypes interaction parameters. The probiotic cocktail consisted of an equal inoculum of *C.pseudodiphtheriticum* and *D. pigrum*, where the normalized inoculum size of each commensal is varied from 10^−1^ to 10^3^. Each treatment course is assumed to last for 5 days and the number of treatment courses varied between 0 (only pretreatment) to 8 (40 days).

### Persistence parameters

Persistence is implemented as a random phenotypic switch between two forms of *S. aureus*: susceptible and persister cells, that are genetically identical. Persisters are the minor subpopulation which are growing slower and showing lower susceptibility to antibiotics relative to the susceptible cells (71). Growth rate of persister *S. aureus* are assumed to be 1/20 of that of susceptible *S. aureus* (e = 0.05) (71), and are assumed to be fully persistent to the application of mupirocin with a relative death rate of 0. Switching rates between two phenotypes are adopted from (71) and (72). Since the mechanisms of how *D. pigrum* and *C. pseudodiphtheriticum* inhibit the growth of *S. aureus* are not exactly clear (39), we assume that persister and susceptible *S. aureus* cells have identical inter- and intra-oligotype interaction terms, i.e., their normalized abundances are equally affected by the normalized abundances of other oligotypes.

### Challenge parameters

Normalized inoculum size and the timing of the *S. aureus* challenge is set to vary between 10^−5^ to 10^3^ and 1 to 120 days following the completion of the antibiotic or probiotic treatment, respectively.

### 4.3. Treatment and challenge simulations

Simulations are implemented as follows, and a brief illustration of the simulation scheme is provided in Fig. 8.

**Figure 8:**
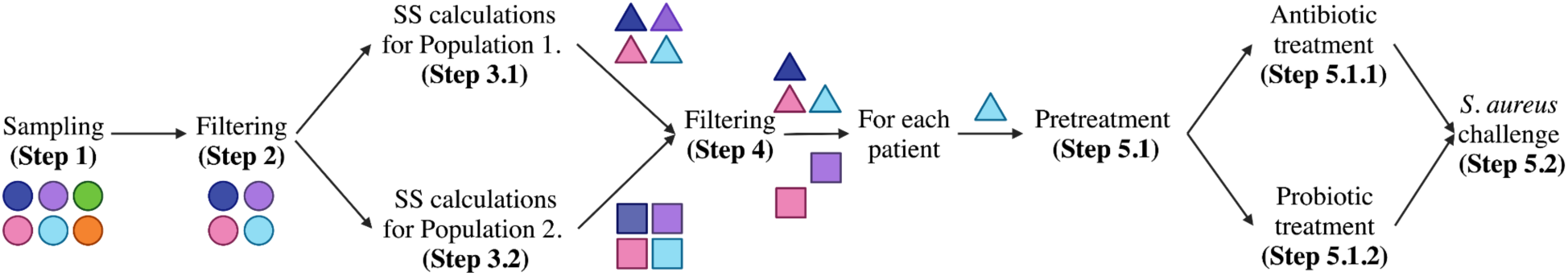
Illustration summarizing the simulation scheme. Color codes represent distinct parameter sets, and the shapes represent the population type, where triangles and squares belong to Population 1 and Population 2, respectively.

1. For each realization of our system, one set of growth and interaction parameters are jointly sampled at random^1^ from the posterior distributions estimated using the 16S rRNA gene sequencing data (44) and the modified version of the MDSINE package (68).
2. Among the sampled parameter sets, ones where *D. pigrum* inhibits the growth of *S. aureus* more than it inhibits the growth of *S. pneumoniae* are chosen.
3. Using these parameter sets, steady state abundances of the dynamical model is obtained for a given set of persistence parameters. Steady states are calculated with the initial conditions reflecting two different patient populations,

3.1. **Population 1:** Initial conditions reflect the normalized steady state abundances with the exception of *D. pigrum* abundance being zero. This condition is based on the inverse association between *S. aureus* carriage and the presence of *D. pigrum* in the nasal microbiome (21–26, 42), and makes the model indifferent to *D. pigrum* and *S. aureus* co-existence.
3.2. **Population 2:** Initial conditions reflect the normalized steady state abundances including the mean value for the *D. pigrum* abundance. This condition imposes *D. pigrum* and *S. aureus* co-existence on the model prior to any treatment.

1. 4. To represent the already colonized patients, parameter sets leading to the coexistence of *S. aureus* and *S. pneumoniae* are chosen. At this step, each parameter set that is chosen for each population represents one distinct patient.
2. 5. For each patient,

5.1. **Pretreatment :** Mupirocin treatment is applied for 5 days. Patients who are still colonized with S. aureus after the pretreatment are chosen for further treatment steps.

5.1.1. **Antibiotic treatment:** Following the pretreatment, mupirocin treatment is continued for a given number of treatment courses, i.e., the abundance of all oligotypes except C1 and C2 is decreased by the killing rate of mupirocin during the simulations.

5.1.2. **Probiotic treatment:** Following the pretreatment, probiotic cocktail is applied for a given number of treatment courses, i.e., the abundance of *Carnobacteriaceae* and C1 are increased by the normalized inoculum size at the time points of probiotic application during the simulations.

Both simulations are continued until the steady-state is reached and the data regarding the microbial abundances are kept for each time step to be later used as the initial conditions for the challenge simulations depending on the time of *S. aureus* inoculation.

5.2. ***S. aureus* challenge:** After the treatment is complete, patients’ nasal cavity is challenged with *S. aureus*, i.e, the normalized abundance of *S. aureus* is increased by the challenge inoculum size at the time point of the challenge during the simulations. Simulations are continued until the new steady-state is reached.

### 4.4. *In vitro* experiments

Human nasal clinical isolates of *C. pseudodiphtheriticum* KPL1989 and *D. pigrum* KPL1914 (30) were used as probiotic cocktails. For the *S. aureus* challenge commercially available methicillin-resistant *S. aureus* (MRSA) USA 300 JE2 strain was used (NARSA Collection, NR-46543, BEI Resources). On the 1st day *C. pseudodiphteriticum* 1989 and *D. pigrum* 1914 were streaked on blood agar plate (Columbia agar + 5% sheep blood, bioMérieux, Marcy-l’Étoile, France) and incubated at 37°C, 5% of CO^2^ for 24 hours. In parallel, nasal swabs (eSwab, Copan, Brescia, Italy) were taken from both nostrils of the donors. 500 ul of the swab liquid amies preservation medium was inoculated to RPMI medium 1640 without phenol red (Life Technologies Corporation, Grand Island, USA) supplemented with 5% tryptic soy broth (TSB) (BD, Franklin Lakes, USA) and with 4 mg/ml mupirocin (Merck, Darmstadt, Germany) for 24h in 37°C, 220rpm.

On the 2nd day, both bacterial strains were transferred separately to 5 ml of RPMI medium with 5% TSB to achieve an optical density (OD) of 0.1. Afterwards both cultures were incubated in 37°C for 6 hours and afterwards bacterial concentration was adjusted to 10^6^ CFU/ml. Nasal swab culture was washed and split equally in two different aliquots (probiotic treatment and control). The probiotic treatment aliquot was supplemented with 1.25 ml of *C. pseudodiphtheriticum* culture and 1.25 ml of *D. pigrum* culture equaling 1.25x 10^6^ CFU each. The negative control was filled up with an RPMI medium with 5% TSB and incubated at 37°C overnight. On the 3rd day both aliquots were challenged with 10^3^ CFU/ml of MRSA and incubated at 37°C. After 24h both aliquots were plated on selective MRSA plates (BBL CHROMagar MRSA II, BD, Franklin Lakes, USA), incubated at 37°C, 5% CO^2^. After 24h incubation CFUs were counted.

## 5. Code and Data Availability

All data and code used in this work is available at https://github.com/burcutepekule/saureusdecolonization.

## Footnotes

1 Each multidimensional vector (which is a part of the joint posterior distribution) has the same probability of being sampled.

