## Supplementary Materials for "Computational and *in vitro* evaluation of probiotic treatments for nasal *Staphylococcus aureus* decolonization"

*These authors contributed equally.

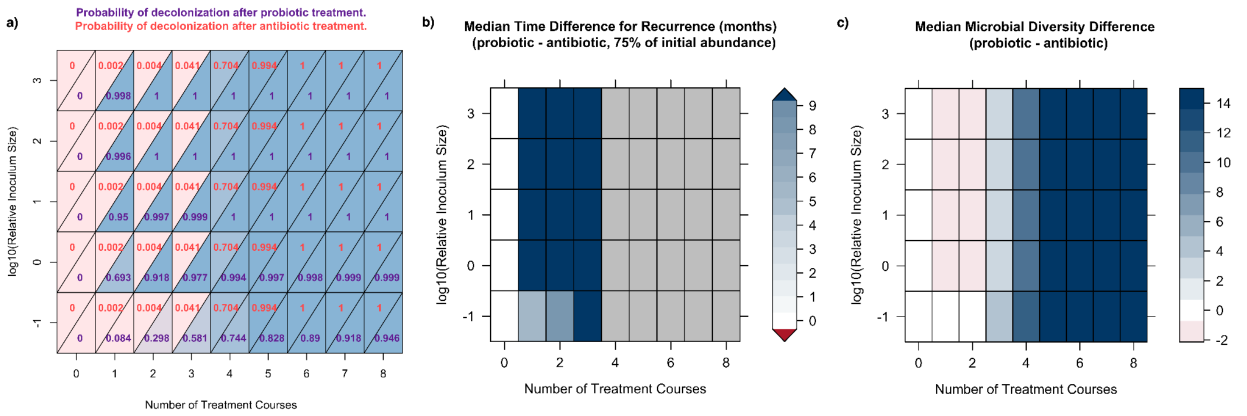


**Figure S1:** Decolonization probability, time to recurrence, and final microbial diversity results for Population 2, with no persisters. **a)** Probability of *S. aureus* decolonization for different number of treatment courses and normalized inoculum sizes. **b)** Median time difference for recurrence between probiotic and antibiotic treatments. **c)** Median microbial diversity difference between probiotic and antibiotic treatments, calculated as the difference of the number of oligotypes still present in the microbiome following treatment.

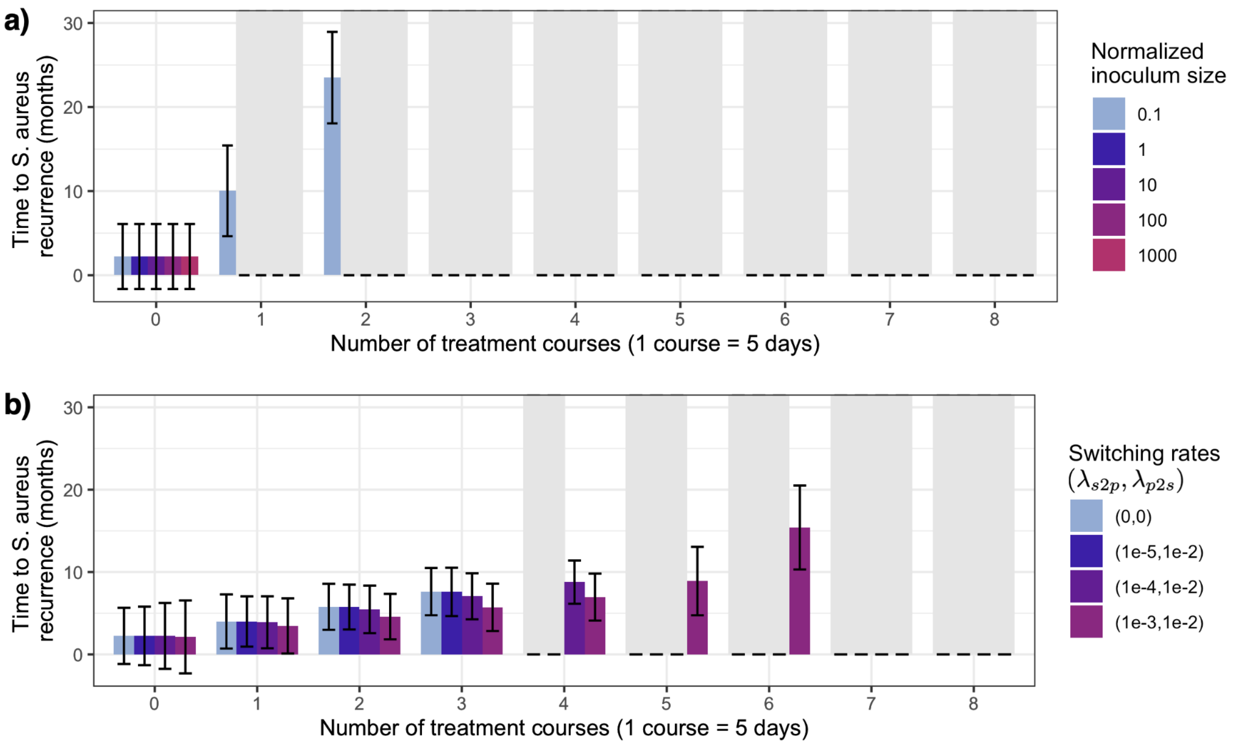


**Figure S2:** Time to *S. aureus* recurrence for Population 2, with different levels of persisters for the antibiotic, and different normalized inoculum sizes for the probiotic treatments. Gray bars represent durations with a mean value longer than 36 months, meaning that no recurrence is observed within the 3 years following treatment. Error bars display +/− the standard error on the mean recurrence time.

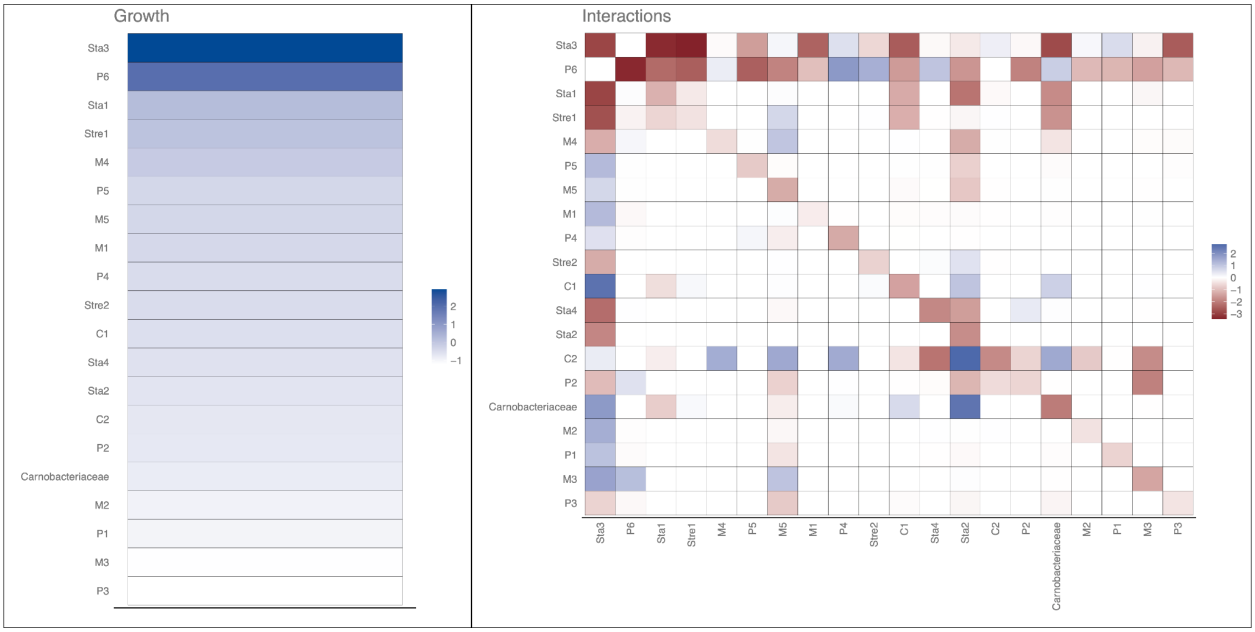


**Figure S3:** Heatmap of the mean values of the growth and interaction parameters estimated by the MDSINE algorithm, values are log10 transformed.

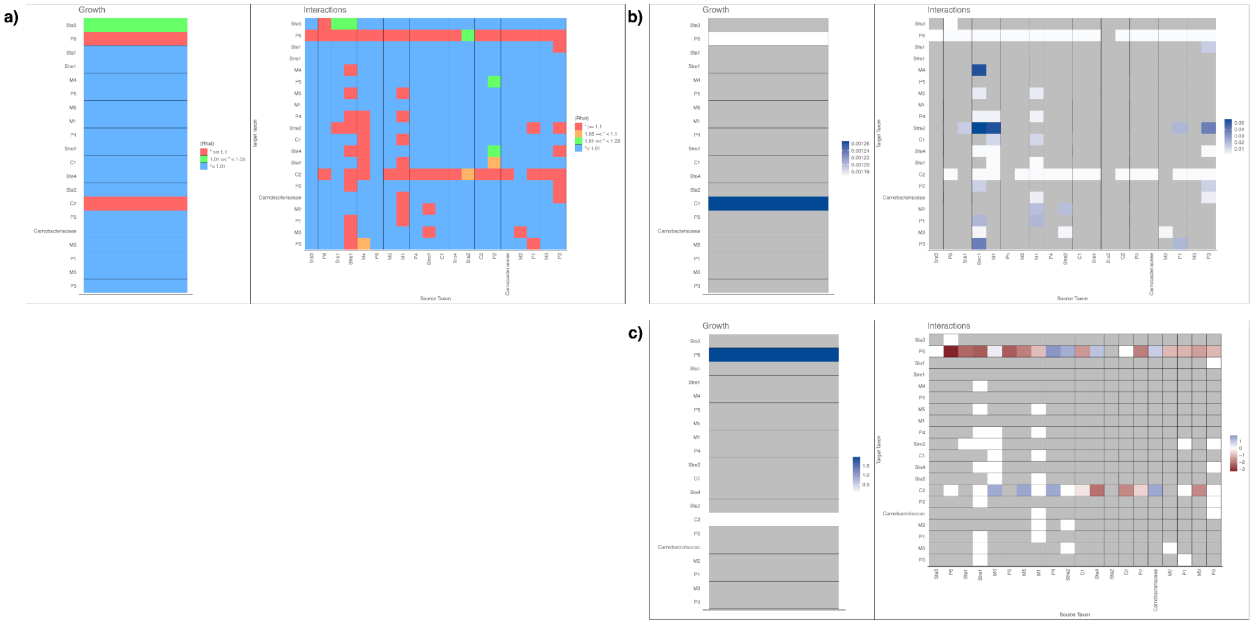


**Figure S4:** Heatmap of the Gelman-Rubin diagnostics for each parameter, where 4 chains are utilized with 25,000 MCMC iterations and 2,500 burn-in iterations. Note that the boxes which are in red have Rhat values exceeding 1.1, which may suggest a potential lack of convergence, indicating that the chains have not mixed well. However, these parameters either related to oligotypes with very low mean abundance, colored boxes contain the information of the mean abundance value for taxon *i* (growth terms), and the minimum of the mean abundance for taxon i and taxon *j* (interaction terms), where *i* and *j* are the row and column indexes, respectively, or have very low values (log10 transformed values of the growth and interaction terms), indicating that although we might lack convergence for these parameters, their impact on the system is minimal.

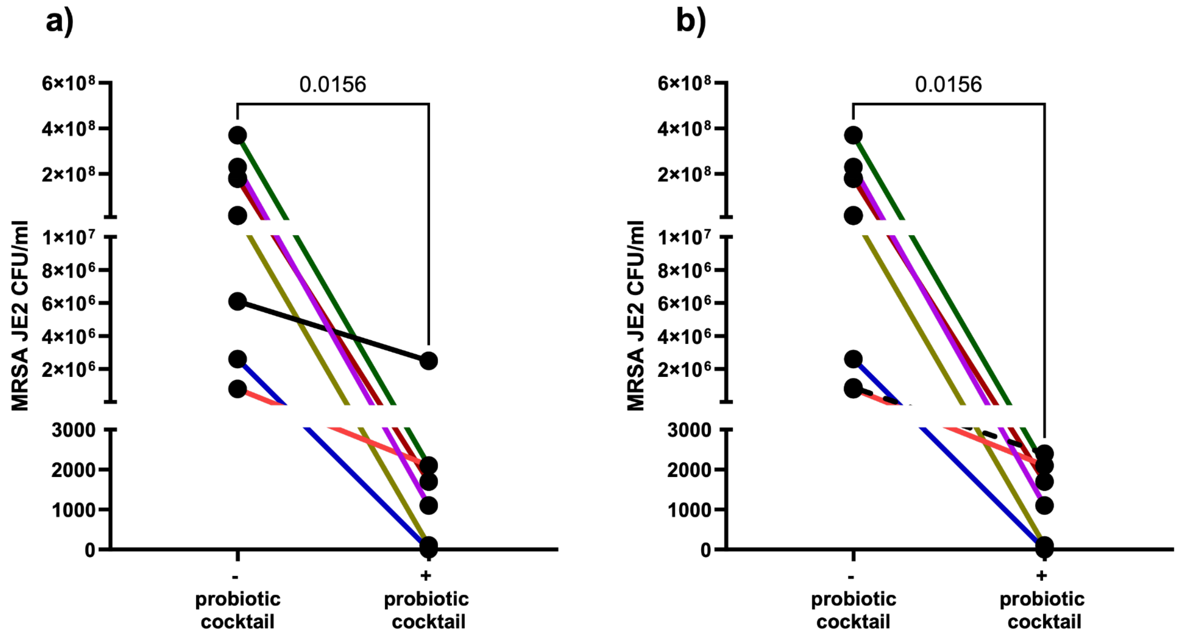


**Figure S5:** Recolonization of MRSA in CFU/ml in the nasal swab liquid aliquots treated with probiotic cocktail containing *D.pigrum* and *C. pseudodiphtheriticum* in a 1:1 ratio versus the untreated aliquots. Separated analysis for one donor, that was repeated (see main text and discussion) - low inhibition of MRSA after probiotic cocktail treatment (**a)**) compared with higher inhibition in the second repetition (**b)**). Individual repeat analysis remains the same as pooled analysis (see Fig. 7 **b)**). For statistical analysis Wilcoxon matched-pairs signed rank test was used.

**Table S1:** Detailed information regarding the growth and interaction parameters, estimated by the MDSINE algorithm. "Source taxon" refers to which taxon or perturbation is affecting the target taxon, and "value" represents the magnitude of the effect (mean of MCMC samples). "Significance" denotes the Bayes factor, and "MCMC std" is the standard deviation across all MCMC samples. Note that the parameters that are intra-species interaction terms and thus forced to take a negative value by the original algorithm do not have a significance value, since the existence of these parameters is enforced. Similarly, the parameters that are conditioned to take a negative/positive value based on the assumptions of this work do not have a significance value.

| parameter_type | source_taxon | target_taxon | value | significance | MCMC_std |
|---|---|---|---|---|---|
| growth_rate | - | Carnobacteriaceae | 0.152 | - | 0.131 |
| growth_rate | - | C1 | 0.288 | - | 0.178 |
| growth_rate | - | C2 | 0.191 | - | 0.179 |
| growth_rate | - | M1 | 0.392 | - | 0.23 |
| growth_rate | - | M2 | 0.114 | - | 0.085 |
| growth_rate | - | M3 | 0.063 | - | 0.048 |
| growth_rate | - | M4 | 0.775 | - | 0.444 |
| growth_rate | - | M5 | 0.427 | - | 0.214 |
| growth_rate | - | P1 | 0.103 | - | 0.086 |
| growth_rate | - | P2 | 0.181 | - | 0.157 |
| growth_rate | - | P3 | 0.061 | - | 0.058 |
| growth_rate | - | P4 | 0.332 | - | 0.131 |
| growth_rate | - | P5 | 0.428 | - | 0.249 |
| growth_rate | - | P6 | 90.999 | - | 0.029 |
| growth_rate | - | Sta1 | 1.62 | - | 0.494 |
| growth_rate | - | Sta2 | 0.218 | - | 0.083 |
| growth_rate | - | Sta3 | 862.137 | - | 317.281 |
| growth_rate | - | Sta4 | 0.253 | - | 0.137 |
| growth_rate | - | Stre1 | 1.141 | - | 0.336 |
| growth_rate | - | Stre2 | 0.33 | - | 0.13 |
| interaction | Carnobacteriaceae | Carnobacteriaceae | -107.541 | - | 23.903 |

| | | | | | |
|---|---|---|---|---|---|
| interaction | Carnobacteriaceae | C1 | 6.377 | - | 6.484 |
| interaction | Carnobacteriaceae | C2 | 32.258 | 0.265 | 64.894 |
| interaction | Carnobacteriaceae | M1 | -0.105 | 0.004 | 2.57 |
| interaction | Carnobacteriaceae | M2 | -0.011 | 0.001 | 0.632 |
| interaction | Carnobacteriaceae | M3 | -0.08 | 0.004 | 1.488 |
| interaction | Carnobacteriaceae | M4 | -2.098 | 0.017 | 28.943 |
| interaction | Carnobacteriaceae | M5 | 0.089 | 0.008 | 9.236 |
| interaction | Carnobacteriaceae | P1 | -0.216 | 0.006 | 3.527 |
| interaction | Carnobacteriaceae | P2 | -0.04 | 0.007 | 10.424 |
| interaction | Carnobacteriaceae | P3 | -0.749 | 0.019 | 5.91 |
| interaction | Carnobacteriaceae | P4 | -0.022 | 0.001 | 0.827 |
| interaction | Carnobacteriaceae | P5 | -0.268 | 0.008 | 10.502 |
| interaction | Carnobacteriaceae | P6 | 7.025 | 0.763 | 765.811 |
| interaction | Carnobacteriaceae | Sta1 | -61.822 | - | 36.895 |
| interaction | Carnobacteriaceae | Sta2 | -0.006 | 0.001 | 0.393 |
| interaction | Carnobacteriaceae | Sta3 | -649.758 | - | 488.301 |
| interaction | Carnobacteriaceae | Sta4 | 0.139 | 0.005 | 5.014 |
| interaction | Carnobacteriaceae | Stre1 | -47.049 | - | 25.037 |
| interaction | Carnobacteriaceae | Stre2 | 0.018 | 0.002 | 0.892 |
| interaction | C1 | Carnobacteriaceae | 4.1 | - | 3.7 |
| interaction | C1 | C1 | -25.918 | - | 9.744 |
| interaction | C1 | C2 | -2.167 | 0.079 | 8.119 |
| interaction | C1 | M1 | -0.238 | 0.008 | 2.971 |
| interaction | C1 | M2 | -0.002 | 0.001 | 0.236 |
| interaction | C1 | M3 | 0 | 0 | 0.074 |
| interaction | C1 | M4 | 0.047 | 0.003 | 2.152 |
| interaction | C1 | M5 | -0.321 | 0.009 | 3.719 |
| interaction | C1 | P1 | -0.001 | 0.001 | 0.251 |
| interaction | C1 | P2 | -0.049 | 0.002 | 1.248 |

| | | | | | |
|---|---|---|---|---|---|
| interaction | C1 | P3 | -0.26 | 0.013 | 2.461 |
| interaction | C1 | P4 | -0.033 | 0.002 | 0.855 |
| interaction | C1 | P5 | 0.106 | 0.004 | 2.693 |
| interaction | C1 | P6 | -34.363 | 0.593 | 567.659 |
| interaction | C1 | Sta1 | -19.606 | - | 12.161 |
| interaction | C1 | Sta2 | -0.012 | 0.002 | 0.351 |
| interaction | C1 | Sta3 | -345.786 | - | 381.717 |
| interaction | C1 | Sta4 | -0.003 | 0.001 | 0.288 |
| interaction | C1 | Stre1 | -16.921 | - | 10.759 |
| interaction | C1 | Stre2 | -0.009 | 0.001 | 0.405 |
| interaction | C2 | Carnobacteriaceae | 0.025 | 0.001 | 1.137 |
| interaction | C2 | C1 | -0.011 | 0.002 | 1.019 |
| interaction | C2 | C2 | -63.736 | - | 35.302 |
| interaction | C2 | M1 | -0.04 | 0.004 | 2.854 |
| interaction | C2 | M2 | 0.042 | 0.002 | 1.542 |
| interaction | C2 | M3 | -0.023 | 0.002 | 0.754 |
| interaction | C2 | M4 | -0.093 | 0.004 | 5.148 |
| interaction | C2 | M5 | -0.037 | 0.003 | 2.336 |
| interaction | C2 | P1 | -0.014 | 0.002 | 0.851 |
| interaction | C2 | P2 | -3.199 | 0.052 | 171.732 |
| interaction | C2 | P3 | 0.01 | 0.002 | 1.329 |
| interaction | C2 | P4 | -0.004 | 0.001 | 0.737 |
| interaction | C2 | P5 | -0.021 | 0.002 | 2.74 |
| interaction | C2 | P6 | 0 | 0 | 0 |
| interaction | C2 | Sta1 | -0.387 | 0.008 | 5.381 |
| interaction | C2 | Sta2 | -0.012 | 0.001 | 0.688 |
| interaction | C2 | Sta3 | 1.402 | 0.057 | 180.766 |
| interaction | C2 | Sta4 | 0.008 | 0.002 | 1.771 |
| interaction | C2 | Stre1 | 0.061 | 0.002 | 2.29 |

| | | | | | |
|---|---|---|---|---|---|
| interaction | C2 | Stre2 | -0.051 | 0.003 | 1.483 |
| interaction | M1 | Carnobacteriaceae | 0 | 0 | 0.009 |
| interaction | M1 | C1 | 0 | 0 | 0.002 |
| interaction | M1 | C2 | -0.003 | 0.002 | 0.076 |
| interaction | M1 | M1 | -1.373 | - | 0.62 |
| interaction | M1 | M2 | 0 | 0 | 0.007 |
| interaction | M1 | M3 | 0 | 0 | 0.007 |
| interaction | M1 | M4 | -0.001 | 0 | 0.076 |
| interaction | M1 | M5 | 0 | 0 | 0.005 |
| interaction | M1 | P1 | 0 | 0 | 0.005 |
| interaction | M1 | P2 | -0.004 | 0.002 | 0.103 |
| interaction | M1 | P3 | 0 | 0 | 0.011 |
| interaction | M1 | P4 | 0 | 0 | 0.008 |
| interaction | M1 | P5 | 0 | 0 | 0.009 |
| interaction | M1 | P6 | -9.647 | 2.258 | 20.392 |
| interaction | M1 | Sta1 | -0.001 | 0 | 0.041 |
| interaction | M1 | Sta2 | 0 | 0 | 0.001 |
| interaction | M1 | Sta3 | -271.276 | 0.244 | 606.168 |
| interaction | M1 | Sta4 | 0 | 0 | 0.011 |
| interaction | M1 | Stre1 | 0 | 0 | 0.029 |
| interaction | M1 | Stre2 | 0 | 0 | 0.006 |
| interaction | M2 | Carnobacteriaceae | 0 | 0 | 0.026 |
| interaction | M2 | C1 | 0.001 | 0 | 0.056 |
| interaction | M2 | C2 | -6.286 | 0.029 | 57.666 |
| interaction | M2 | M1 | 0.001 | 0 | 0.089 |
| interaction | M2 | M2 | -2.383 | - | 1.383 |
| interaction | M2 | M3 | 0 | 0 | 0 |
| interaction | M2 | M4 | -0.001 | 0 | 0.062 |
| interaction | M2 | M5 | 0 | 0 | 0.052 |

| | | | | | |
|---|---|---|---|---|---|
| interaction | M2 | P1 | 0 | 0 | 0.013 |
| interaction | M2 | P2 | -0.001 | 0 | 0.056 |
| interaction | M2 | P3 | 0 | 0 | 0.046 |
| interaction | M2 | P4 | 0.004 | 0.001 | 0.245 |
| interaction | M2 | P5 | -0.006 | 0.001 | 0.178 |
| interaction | M2 | P6 | -11.431 | 2.898 | 134.27 |
| interaction | M2 | Sta1 | 0 | 0 | 0.033 |
| interaction | M2 | Sta2 | 0.001 | 0.001 | 0.071 |
| interaction | M2 | Sta3 | 0.598 | 0.054 | 181.796 |
| interaction | M2 | Sta4 | 0 | 0 | 0.038 |
| interaction | M2 | Stre1 | 0 | 0 | 0.057 |
| interaction | M2 | Stre2 | 0 | 0 | 0.013 |
| interaction | M3 | Carnobacteriaceae | -0.069 | 0.003 | 2.321 |
| interaction | M3 | C1 | 0.05 | 0.003 | 1.878 |
| interaction | M3 | C2 | -58.556 | 0.15 | 164.047 |
| interaction | M3 | M1 | -0.095 | 0.004 | 2.571 |
| interaction | M3 | M2 | -0.01 | 0.001 | 0.874 |
| interaction | M3 | M3 | -23.55 | - | 9.256 |
| interaction | M3 | M4 | -0.24 | 0.007 | 8.423 |
| interaction | M3 | M5 | -0.038 | 0.003 | 2.532 |
| interaction | M3 | P1 | -0.018 | 0.002 | 1.286 |
| interaction | M3 | P2 | -88.933 | 0.115 | 309.157 |
| interaction | M3 | P3 | 0.01 | 0.002 | 0.862 |
| interaction | M3 | P4 | 0.009 | 0.001 | 0.508 |
| interaction | M3 | P5 | -0.015 | 0.004 | 2.683 |
| interaction | M3 | P6 | -27.749 | 0.615 | 650.46 |
| interaction | M3 | Sta1 | -0.626 | 0.01 | 7.672 |
| interaction | M3 | Sta2 | 0.011 | 0.001 | 0.549 |
| interaction | M3 | Sta3 | -0.988 | 0.054 | 188.258 |

| | | | | | |
|---|---|---|---|---|---|
| interaction | M3 | Sta4 | -0.038 | 0.002 | 2.652 |
| interaction | M3 | Stre1 | 0.028 | 0.003 | 2.017 |
| interaction | M3 | Stre2 | 0.006 | 0.001 | 0.685 |
| interaction | M4 | Carnobacteriaceae | 0 | 0 | 0.027 |
| interaction | M4 | C1 | 0 | 0 | 0 |
| interaction | M4 | C2 | 28.874 | 0.059 | 234.939 |
| interaction | M4 | M1 | -0.01 | 0.003 | 0.199 |
| interaction | M4 | M2 | -0.001 | 0.001 | 0.037 |
| interaction | M4 | M3 | 0 | 0 | 0.021 |
| interaction | M4 | M4 | -3.11 | - | 1.415 |
| interaction | M4 | M5 | 0 | 0 | 0.019 |
| interaction | M4 | P1 | -0.005 | 0.002 | 0.121 |
| interaction | M4 | P2 | 0.001 | 0 | 0.055 |
| interaction | M4 | P3 | 0 | 0 | 0.009 |
| interaction | M4 | P4 | 0 | 0 | 0 |
| interaction | M4 | P5 | 0 | 0 | 0.035 |
| interaction | M4 | P6 | 1.551 | 2.125 | 4.024 |
| interaction | M4 | Sta1 | 0 | 0 | 0.009 |
| interaction | M4 | Sta2 | 0 | 0 | 0.005 |
| interaction | M4 | Sta3 | -0.336 | 0.055 | 186.867 |
| interaction | M4 | Sta4 | 0 | 0 | 0.009 |
| interaction | M4 | Stre1 | -0.001 | 0 | 0.047 |
| interaction | M4 | Stre2 | 0 | 0 | 0 |
| interaction | M5 | Carnobacteriaceae | -1.328 | 0.012 | 17.237 |
| interaction | M5 | C1 | 0.148 | 0.005 | 6.07 |
| interaction | M5 | C2 | 33.712 | 0.068 | 180.838 |
| interaction | M5 | M1 | 0.189 | 0.005 | 3.098 |
| interaction | M5 | M2 | -0.557 | 0.008 | 8.825 |
| interaction | M5 | M3 | 10.956 | 0.042 | 63.195 |

| | | | | | |
|---|---|---|---|---|---|
| interaction | M5 | M4 | 9.893 | 0.044 | 54.06 |
| interaction | M5 | M5 | -19.042 | - | 3.069 |
| interaction | M5 | P1 | -2.11 | 0.016 | 24.644 |
| interaction | M5 | P2 | -4.793 | 0.017 | 61.372 |
| interaction | M5 | P3 | -6.075 | 0.032 | 38.959 |
| interaction | M5 | P4 | -1.288 | 0.013 | 17.648 |
| interaction | M5 | P5 | -0.237 | 0.007 | 10.444 |
| interaction | M5 | P6 | -82.191 | 0.707 | 648.01 |
| interaction | M5 | Sta1 | 0.244 | 0.008 | 11.23 |
| interaction | M5 | Sta2 | -0.016 | 0.002 | 1.207 |
| interaction | M5 | Sta3 | 0.717 | 0.055 | 189.112 |
| interaction | M5 | Sta4 | -0.485 | 0.016 | 14.387 |
| interaction | M5 | Stre1 | 4.736 | 0.094 | 16.418 |
| interaction | M5 | Stre2 | 0.064 | 0.003 | 1.312 |
| interaction | P1 | Carnobacteriaceae | 0 | 0 | 0.039 |
| interaction | P1 | C1 | -0.001 | 0 | 0.051 |
| interaction | P1 | C2 | -0.039 | 0.011 | 0.379 |
| interaction | P1 | M1 | -0.004 | 0.001 | 0.162 |
| interaction | P1 | M2 | -0.001 | 0 | 0.051 |
| interaction | P1 | M3 | -0.001 | 0 | 0.037 |
| interaction | P1 | M4 | 0.002 | 0.001 | 0.356 |
| interaction | P1 | M5 | 0 | 0 | 0.026 |
| interaction | P1 | P1 | -4.399 | - | 1.701 |
| interaction | P1 | P2 | 0 | 0 | 0.048 |
| interaction | P1 | P3 | 0 | 0 | 0.032 |
| interaction | P1 | P4 | 0 | 0 | 0.056 |
| interaction | P1 | P5 | 0.004 | 0.001 | 0.175 |
| interaction | P1 | P6 | -13.32 | 0.621 | 849.805 |
| interaction | P1 | Sta1 | 0.004 | 0.001 | 0.156 |

| | | | | | |
|---|---|---|---|---|---|
| interaction | P1 | Sta2 | 0.001 | 0 | 0.06 |
| interaction | P1 | Sta3 | 3.703 | 0.053 | 169.922 |
| interaction | P1 | Sta4 | 0 | 0 | 0.034 |
| interaction | P1 | Stre1 | 0.001 | 0 | 0.074 |
| interaction | P1 | Stre2 | 0 | 0 | 0.018 |
| interaction | P2 | Carnobacteriaceae | 0 | 0 | 0.112 |
| interaction | P2 | C1 | 0 | 0 | 0.063 |
| interaction | P2 | C2 | -4.291 | 0.043 | 25.264 |
| interaction | P2 | M1 | 0 | 0 | 0.03 |
| interaction | P2 | M2 | 0 | 0 | 0.019 |
| interaction | P2 | M3 | 0 | 0 | 0.035 |
| interaction | P2 | M4 | 0 | 0 | 0.073 |
| interaction | P2 | M5 | 0 | 0 | 0.019 |
| interaction | P2 | P1 | 0 | 0 | 0.029 |
| interaction | P2 | P2 | -4.102 | - | 2.147 |
| interaction | P2 | P3 | 0 | 0 | 0.027 |
| interaction | P2 | P4 | 0.001 | 0 | 0.088 |
| interaction | P2 | P5 | 0 | 0 | 0.022 |
| interaction | P2 | P6 | -85.559 | 4.108 | 69.595 |
| interaction | P2 | Sta1 | 0 | 0 | 0.042 |
| interaction | P2 | Sta2 | 0 | 0 | 0.028 |
| interaction | P2 | Sta3 | -0.481 | 0.053 | 177.357 |
| interaction | P2 | Sta4 | 1.898 | 0.254 | 3.918 |
| interaction | P2 | Stre1 | 0 | 0 | 0.065 |
| interaction | P2 | Stre2 | 0 | 0 | 0.031 |
| interaction | P3 | Carnobacteriaceae | 0 | 0 | 0.011 |
| interaction | P3 | C1 | 0.001 | 0 | 0.059 |
| interaction | P3 | C2 | 0 | 0 | 0.044 |
| interaction | P3 | M1 | 0 | 0 | 0.044 |

| | | | | | |
|---|---|---|---|---|---|
| interaction | P3 | M2 | 0 | 0 | 0.058 |
| interaction | P3 | M3 | 0.001 | 0.001 | 0.041 |
| interaction | P3 | M4 | -0.301 | 0.006 | 6.912 |
| interaction | P3 | M5 | -0.001 | 0 | 0.069 |
| interaction | P3 | P1 | 0 | 0 | 0.013 |
| interaction | P3 | P2 | 0 | 0 | 0.019 |
| interaction | P3 | P3 | -2.101 | - | 1.172 |
| interaction | P3 | P4 | -0.001 | 0 | 0.048 |
| interaction | P3 | P5 | -0.163 | 0.004 | 4.071 |
| interaction | P3 | P6 | -12.101 | 1.199 | 540.621 |
| interaction | P3 | Sta1 | 0 | 0 | 0.016 |
| interaction | P3 | Sta2 | -0.001 | 0 | 0.031 |
| interaction | P3 | Sta3 | -349.329 | 0.238 | 819.222 |
| interaction | P3 | Sta4 | 0 | 0 | 0.014 |
| interaction | P3 | Stre1 | 0 | 0 | 0.004 |
| interaction | P3 | Stre2 | 0 | 0 | 0 |
| interaction | P4 | Carnobacteriaceae | 0.423 | 0.017 | 3.502 |
| interaction | P4 | C1 | 0.002 | 0.001 | 0.283 |
| interaction | P4 | C2 | 31.083 | 0.148 | 84.254 |
| interaction | P4 | M1 | -0.047 | 0.002 | 1.218 |
| interaction | P4 | M2 | -0.026 | 0.002 | 0.787 |
| interaction | P4 | M3 | -0.004 | 0.001 | 0.26 |
| interaction | P4 | M4 | 0.004 | 0.002 | 0.992 |
| interaction | P4 | M5 | 0.013 | 0.001 | 0.695 |
| interaction | P4 | P1 | -0.01 | 0.001 | 0.476 |
| interaction | P4 | P2 | 0.047 | 0.002 | 1.178 |
| interaction | P4 | P3 | 0.016 | 0.002 | 0.573 |
| interaction | P4 | P4 | -19.719 | - | 7.815 |
| interaction | P4 | P5 | -0.013 | 0.001 | 0.735 |

| | | | | | |
|---|---|---|---|---|---|
| interaction | P4 | P6 | 65.352 | 2.042 | 492.992 |
| interaction | P4 | Sta1 | -0.011 | 0.001 | 0.677 |
| interaction | P4 | Sta2 | 0.004 | 0.001 | 0.205 |
| interaction | P4 | Sta3 | 3.181 | 0.054 | 183.561 |
| interaction | P4 | Sta4 | 0.013 | 0.001 | 0.441 |
| interaction | P4 | Stre1 | -0.024 | 0.002 | 0.942 |
| interaction | P4 | Stre2 | 0 | 0.001 | 0.262 |
| interaction | P5 | Carnobacteriaceae | -0.003 | 0.001 | 0.181 |
| interaction | P5 | C1 | 0 | 0 | 0.068 |
| interaction | P5 | C2 | -0.004 | 0.001 | 0.148 |
| interaction | P5 | M1 | -0.001 | 0.001 | 0.18 |
| interaction | P5 | M2 | -0.001 | 0 | 0.088 |
| interaction | P5 | M3 | -0.001 | 0 | 0.068 |
| interaction | P5 | M4 | -0.089 | 0.003 | 2.233 |
| interaction | P5 | M5 | 0 | 0 | 0.082 |
| interaction | P5 | P1 | -0.009 | 0.001 | 0.282 |
| interaction | P5 | P2 | -0.002 | 0 | 0.12 |
| interaction | P5 | P3 | -0.002 | 0.001 | 0.138 |
| interaction | P5 | P4 | 0.73 | 0.011 | 8.788 |
| interaction | P5 | P5 | -6.146 | - | 4.146 |
| interaction | P5 | P6 | -296.796 | 0.811 | 1019.018 |
| interaction | P5 | Sta1 | 0.006 | 0.001 | 0.312 |
| interaction | P5 | Sta2 | -0.005 | 0.001 | 0.236 |
| interaction | P5 | Sta3 | -31.472 | 0.068 | 244.617 |
| interaction | P5 | Sta4 | 0 | 0 | 0.031 |
| interaction | P5 | Stre1 | 0.004 | 0.001 | 0.24 |
| interaction | P5 | Stre2 | 0 | 0 | 0.121 |
| interaction | P6 | Carnobacteriaceae | -0.025 | 0.003 | 2.265 |
| interaction | P6 | C1 | 0.003 | 0.002 | 1.039 |

| | | | | | |
|---|---|---|---|---|---|
| interaction | P6 | C2 | 0 | 0 | 0 |
| interaction | P6 | M1 | -0.493 | 0.008 | 7.016 |
| interaction | P6 | M2 | -0.113 | 0.004 | 2.341 |
| interaction | P6 | M3 | 13.426 | 0.042 | 128.019 |
| interaction | P6 | M4 | 0.691 | 0.01 | 9.459 |
| interaction | P6 | M5 | 0.097 | 0.004 | 2.693 |
| interaction | P6 | P1 | -0.282 | 0.005 | 4.734 |
| interaction | P6 | P2 | 2.918 | 0.019 | 26.94 |
| interaction | P6 | P3 | -0.457 | 0.008 | 6.884 |
| interaction | P6 | P4 | -0.138 | 0.004 | 2.893 |
| interaction | P6 | P5 | -0.096 | 0.003 | 3.199 |
| interaction | P6 | P6 | -2172.93 | - | 2.498 |
| interaction | P6 | Sta1 | 0.239 | 0.005 | 4.728 |
| interaction | P6 | Sta2 | 0.002 | 0.001 | 0.335 |
| interaction | P6 | Sta3 | 0 | 0 | 0 |
| interaction | P6 | Sta4 | 0.073 | 0.004 | 2.063 |
| interaction | P6 | Stre1 | -0.906 | 0.013 | 9.245 |
| interaction | P6 | Stre2 | 0.027 | 0.002 | 1.368 |
| interaction | Sta1 | Carnobacteriaceae | -5.576 | - | 2.799 |
| interaction | Sta1 | C1 | -2.847 | - | 2.982 |
| interaction | Sta1 | C2 | -1.296 | 0.161 | 3.581 |
| interaction | Sta1 | M1 | 0 | 0 | 0.112 |
| interaction | Sta1 | M2 | 0 | 0 | 0.029 |
| interaction | Sta1 | M3 | 0 | 0 | 0 |
| interaction | Sta1 | M4 | 0.001 | 0 | 0.079 |
| interaction | Sta1 | M5 | -0.001 | 0 | 0.104 |
| interaction | Sta1 | P1 | 0 | 0 | 0.042 |
| interaction | Sta1 | P2 | -0.001 | 0 | 0.093 |
| interaction | Sta1 | P3 | -0.001 | 0 | 0.07 |

| | | | | | |
|---|---|---|---|---|---|
| interaction | Sta1 | P4 | -0.011 | 0.001 | 0.502 |
| interaction | Sta1 | P5 | 0.004 | 0.001 | 0.165 |
| interaction | Sta1 | P6 | -202.022 | 2.161 | 360.986 |
| interaction | Sta1 | Sta1 | -15.112 | - | 4.089 |
| interaction | Sta1 | Sta2 | -0.001 | 0 | 0.049 |
| interaction | Sta1 | Sta3 | -2050.57 | - | 857.857 |
| interaction | Sta1 | Sta4 | -0.001 | 0 | 0.046 |
| interaction | Sta1 | Stre1 | -4.11 | - | 2.888 |
| interaction | Sta1 | Stre2 | 0 | 0 | 0.025 |
| interaction | Sta2 | Carnobacteriaceae | 340.804 | 32.186 | 103.19 |
| interaction | Sta2 | C1 | 10.451 | 0.06 | 47.892 |
| interaction | Sta2 | C2 | 571.047 | 1.867 | 458.123 |
| interaction | Sta2 | M1 | -0.242 | 0.008 | 8.439 |
| interaction | Sta2 | M2 | -0.087 | 0.004 | 3.364 |
| interaction | Sta2 | M3 | 0.118 | 0.004 | 3.179 |
| interaction | Sta2 | M4 | -18.929 | 0.044 | 123.058 |
| interaction | Sta2 | M5 | -6.761 | 0.027 | 60.87 |
| interaction | Sta2 | P1 | -0.391 | 0.007 | 6.816 |
| interaction | Sta2 | P2 | -13.338 | 0.039 | 85.848 |
| interaction | Sta2 | P3 | -0.658 | 0.009 | 9.47 |
| interaction | Sta2 | P4 | -0.044 | 0.003 | 3.539 |
| interaction | Sta2 | P5 | -4.877 | 0.024 | 52.013 |
| interaction | Sta2 | P6 | -42.015 | 0.568 | 563.099 |
| interaction | Sta2 | Sta1 | -141.246 | 0.233 | 334.02 |
| interaction | Sta2 | Sta2 | -57.364 | - | 28.569 |
| interaction | Sta2 | Sta3 | -1.685 | 0.053 | 191.444 |
| interaction | Sta2 | Sta4 | -31.697 | 0.103 | 110.528 |
| interaction | Sta2 | Stre1 | -0.609 | 0.009 | 13.628 |
| interaction | Sta2 | Stre2 | 2.773 | 0.025 | 19.74 |

| | | | | | |
|---|---|---|---|---|---|
| interaction | Sta3 | Carnobacteriaceae | 62.54 | - | 654.472 |
| interaction | Sta3 | C1 | 362.482 | - | 620.82 |
| interaction | Sta3 | C2 | 1.735 | 0.036 | 109.604 |
| interaction | Sta3 | M1 | 15.877 | 0.049 | 161.596 |
| interaction | Sta3 | M2 | 27.015 | 0.052 | 163.746 |
| interaction | Sta3 | M3 | 41.787 | 0.078 | 173.716 |
| interaction | Sta3 | M4 | -17.562 | 0.052 | 176.494 |
| interaction | Sta3 | M5 | 4.784 | 0.043 | 132.891 |
| interaction | Sta3 | P1 | 11.462 | 0.037 | 117.442 |
| interaction | Sta3 | P2 | -10.666 | 0.047 | 155.831 |
| interaction | Sta3 | P3 | -4.415 | 0.04 | 121.304 |
| interaction | Sta3 | P4 | 3.062 | 0.041 | 121.242 |
| interaction | Sta3 | P5 | 15.971 | 0.05 | 162.641 |
| interaction | Sta3 | P6 | 0 | 0 | 0 |
| interaction | Sta3 | Sta1 | -816.516 | - | 532.179 |
| interaction | Sta3 | Sta2 | -72.039 | 0.123 | 232.344 |
| interaction | Sta3 | Sta3 | -760.308 | - | 683.472 |
| interaction | Sta3 | Sta4 | -177.311 | 0.24 | 403.641 |
| interaction | Sta3 | Stre1 | -529.112 | - | 406.773 |
| interaction | Sta3 | Stre2 | -18.877 | 0.046 | 143.835 |
| interaction | Sta4 | Carnobacteriaceae | 0.007 | 0.001 | 0.713 |
| interaction | Sta4 | C1 | 0.006 | 0.002 | 0.68 |
| interaction | Sta4 | C2 | -140.868 | 0.46 | 271.932 |
| interaction | Sta4 | M1 | -0.202 | 0.005 | 3.666 |
| interaction | Sta4 | M2 | 0.063 | 0.003 | 1.554 |
| interaction | Sta4 | M3 | -0.001 | 0 | 0.193 |
| interaction | Sta4 | M4 | 0.006 | 0.002 | 1.399 |
| interaction | Sta4 | M5 | -0.033 | 0.002 | 1.424 |
| interaction | Sta4 | P1 | -0.037 | 0.002 | 1.176 |

| | | | | | |
|---|---|---|---|---|---|
| interaction | Sta4 | P2 | -0.232 | 0.006 | 3.502 |
| interaction | Sta4 | P3 | -0.018 | 0.001 | 0.913 |
| interaction | Sta4 | P4 | 0.013 | 0.002 | 1.353 |
| interaction | Sta4 | P5 | -0.096 | 0.003 | 2.192 |
| interaction | Sta4 | P6 | 10.374 | 2.124 | 775.646 |
| interaction | Sta4 | Sta1 | -0.005 | 0.002 | 1.284 |
| interaction | Sta4 | Sta2 | 0.012 | 0.002 | 0.51 |
| interaction | Sta4 | Sta3 | -0.442 | 0.053 | 181.958 |
| interaction | Sta4 | Sta4 | -68.818 | - | 15.383 |
| interaction | Sta4 | Stre1 | -0.065 | 0.003 | 2.291 |
| interaction | Sta4 | Stre2 | 0.256 | 0.007 | 3.368 |
| interaction | Stre1 | Carnobacteriaceae | 0.511 | - | 0.548 |
| interaction | Stre1 | C1 | 0.558 | - | 0.576 |
| interaction | Stre1 | C2 | 0 | 0 | 0.016 |
| interaction | Stre1 | M1 | 0 | 0 | 0.014 |
| interaction | Stre1 | M2 | 0 | 0 | 0.01 |
| interaction | Stre1 | M3 | 0 | 0 | 0 |
| interaction | Stre1 | M4 | 0 | 0 | 0.02 |
| interaction | Stre1 | M5 | 0 | 0 | 0.031 |
| interaction | Stre1 | P1 | 0 | 0 | 0.004 |
| interaction | Stre1 | P2 | 0 | 0 | 0.02 |
| interaction | Stre1 | P3 | 0 | 0 | 0.009 |
| interaction | Stre1 | P4 | 0 | 0 | 0.012 |
| interaction | Stre1 | P5 | 0.001 | 0 | 0.034 |
| interaction | Stre1 | P6 | -315.206 | 11.806 | 142.432 |
| interaction | Stre1 | Sta1 | -1.671 | - | 1.015 |
| interaction | Stre1 | Sta2 | 0 | 0 | 0.012 |
| interaction | Stre1 | Sta3 | -2657.16 | - | 837.518 |
| interaction | Stre1 | Sta4 | 0 | 0 | 0.005 |

| | | | | | |
|---|---|---|---|---|---|
| interaction | Stre1 | Stre1 | -2.325 | - | 0.905 |
| interaction | Stre1 | Stre2 | 0 | 0 | 0 |
| interaction | Stre2 | Carnobacteriaceae | 0 | 0 | 0.023 |
| interaction | Stre2 | C1 | 0.338 | 0.023 | 2.39 |
| interaction | Stre2 | C2 | 0.014 | 0.004 | 0.727 |
| interaction | Stre2 | M1 | 0 | 0 | 0.055 |
| interaction | Stre2 | M2 | 0 | 0 | 0.024 |
| interaction | Stre2 | M3 | 0 | 0 | 0.058 |
| interaction | Stre2 | M4 | 0.022 | 0.002 | 0.949 |
| interaction | Stre2 | M5 | -0.001 | 0.001 | 0.079 |
| interaction | Stre2 | P1 | -0.004 | 0.001 | 0.161 |
| interaction | Stre2 | P2 | -0.003 | 0 | 0.186 |
| interaction | Stre2 | P3 | -0.008 | 0.002 | 0.196 |
| interaction | Stre2 | P4 | -0.002 | 0.001 | 0.103 |
| interaction | Stre2 | P5 | -0.002 | 0.001 | 0.53 |
| interaction | Stre2 | P6 | 24.941 | 1.272 | 820.246 |
| interaction | Stre2 | Sta1 | -0.002 | 0.001 | 0.257 |
| interaction | Stre2 | Sta2 | -0.002 | 0.001 | 0.073 |
| interaction | Stre2 | Sta3 | -3.803 | 0.056 | 184.82 |
| interaction | Stre2 | Sta4 | 0.001 | 0.001 | 0.074 |
| interaction | Stre2 | Stre1 | 0.066 | 0.005 | 0.998 |
| interaction | Stre2 | Stre2 | -4.614 | - | 1.299 |
